# Lipid availability influences ferroptosis sensitivity in cancer cells by regulating polyunsaturated fatty acid trafficking

**DOI:** 10.1101/2024.05.06.592780

**Authors:** Kelly H. Sokol, Cameron J. Lee, Thomas J. Rogers, Althea Waldhart, Abigail E. Ellis, Sahithi Madireddy, Samuel R. Daniels, Xinyu Ye, Mary Olesnavich, Amy Johnson, Benjamin R. Furness, Ryan D. Sheldon, Evan C. Lien

## Abstract

Ferroptosis is a form of cell death caused by lipid peroxidation that is emerging as a target for cancer therapy, highlighting the need to identify factors that govern ferroptosis susceptibility. Lipid peroxidation occurs primarily on phospholipids containing polyunsaturated fatty acids (PUFAs). Here, we show that even though extracellular lipid limitation reduces cellular PUFA levels, lipid-starved cancer cells are paradoxically more sensitive to ferroptosis. Using mass spectrometry-based lipidomics with stable isotope fatty acid labeling, we show that lipid limitation induces a fatty acid trafficking pathway in which PUFAs are liberated from triglycerides to synthesize highly unsaturated PUFAs such as arachidonic acid and adrenic acid. These PUFAs then accumulate in phospholipids, particularly ether phospholipids, to promote ferroptosis sensitivity. Therefore, PUFA levels within cancer cells do not necessarily correlate with ferroptosis susceptibility. Rather, how cancer cells respond to extracellular lipid levels by trafficking PUFAs into proper phospholipid pools dictates their sensitivity to ferroptosis.

## INTRODUCTION

Ferroptosis is a non-apoptotic form of cell death that is triggered by oxidative damage to cellular phospholipids^1,2^. The peroxidation of lipids occurs in an iron-dependent manner, leading to the formation of toxic lipid hydroperoxides in cellular membranes. Under normal conditions, cells have a lipid peroxide repair system, in which lipid peroxides are reduced to non-toxic lipid alcohols by glutathione (GSH)-dependent lipid hydroperoxidase glutathione peroxidase 4 (GPX4). Impairing lipid peroxide repair, either through direct inhibition of GPX4 or blocking the production of antioxidants such as GSH and coenzyme Q_10_, results in the cellular accumulation of lipid peroxides that induce ferroptosis^1,2^. Because ferroptosis has been implicated in several pathological conditions, including cancer, ischemia-reperfusion injury, and neurodegenerative diseases, manipulating ferroptosis induction is being explored as a potential strategy for new therapies^1,2^.

Lipid peroxidation occurs primarily within phospholipids containing polyunsaturated fatty acids (PUFAs), which are more susceptible to oxidation because they have multiple double bonds with bis-allylic hydrogen atoms^3,4^. In particular, arachidonic acid (20:4(n-6)) and adrenic acid (22:4(n-6)) have been implicated in ferroptosis induction in some cell types, and supplementing cells with exogenous PUFAs such as 20:4(n-6) can sensitize cells to ferroptosis through their incorporation into phospholipids^5–9^. A recent study also suggests that phospholipids with two PUFA chains are particularly potent at promoting ferroptosis^4^. In contrast, saturated fatty acids (SFAs) and monounsaturated fatty acids (MUFAs) are significantly less susceptible to oxidation^3^. Moreover, providing cells with MUFAs inhibits ferroptosis by displacing PUFA-containing phospholipids from the plasma membrane^10^. These results have led to the current paradigm that increased cellular PUFA levels promote ferroptosis sensitivity, whereas higher cellular MUFA levels confer ferroptosis resistance.

We previously demonstrated that extracellular lipid limitation alters the relative composition of SFAs, MUFAs, and PUFAs in cancer cells. Lipid-starved cells maintain SFA and MUFA levels by stimulating *de novo* fatty acid synthesis, and even have higher levels of MUFAs such as palmitoleate (16:1(n-7)) through activation of stearoyl-CoA desaturase (SCD)^11,12^. In contrast, because PUFAs are essential fatty acids that must be obtained from the environment, PUFA levels drop in lipid-starved cancer cells^11^. Because PUFAs are required for ferroptosis and MUFAs confer ferroptosis resistance^1,3,10^, we hypothesized that extracellular lipid limitation would cause MUFA-rich, PUFA-deficient cancer cells to become resistant to ferroptosis. To our surprise, we found that lipid-starved cancer cells are instead more sensitive to ferroptosis induction by the GPX4 inhibitor RSL3. Using a combination of mass spectrometry-based fatty acid and lipidomics analyses and stable isotope fatty acid labeling, we demonstrate that lipid-starved cancer cells activate a PUFA trafficking pathway in which PUFAs are liberated from triglycerides and used to synthesize long chain, highly unsaturated fatty acids such as 20:4(n-6) and 22:4(n-6). These PUFAs are then incorporated into phospholipids, in particular ether phospholipids, to promote ferroptosis sensitivity. These results highlight that it is not the overall cellular levels of PUFAs, but rather the intracellular trafficking of highly unsaturated PUFAs into the proper phospholipid pools, that dictate ferroptosis sensitivity in cancer cells.

## RESULTS

### Extracellular lipid limitation sensitizes human cancer cells to ferroptosis

We previously developed a method to culture cells in lipid-depleted media^13^. Under these conditions, we showed that cancer cells up-regulate *de novo* fatty acid synthesis and the production of MUFAs via stearoyl-CoA desaturase (SCD)^11,12^. Consistently, by using gas chromatography-mass spectrometry (GC-MS) to profile the cellular fatty acid composition of a panel of human cancer cell lines (A549 lung cancer cells, HeLa cervical cancer cells, Panc1 pancreatic cancer cells, H1299 lung cancer cells, and MIA PaCa-2 pancreatic cancer cells) grown in lipid-replete versus lipid-depleted culture media, we found that lipid-starved cancer cells had elevated levels of the MUFA palmitoleic acid (16:1(n-7); Fig. 1A). Extracellular lipid limitation also increased the ratios of the MUFAs 16:1(n-7) and oleic acid (18:1(n-9)) relative to their respective SFA precursors, palmitate (16:0) and oleate (18:0), which reflects increased SCD activity (Fig. 1A). In contrast, lipid-starved cancer cells had lower levels of PUFAs (Fig. 1A), which are essential fatty acids that cannot be synthesized *de novo* by mammalian cells and must be obtained from the environment.

**Figure 1.**
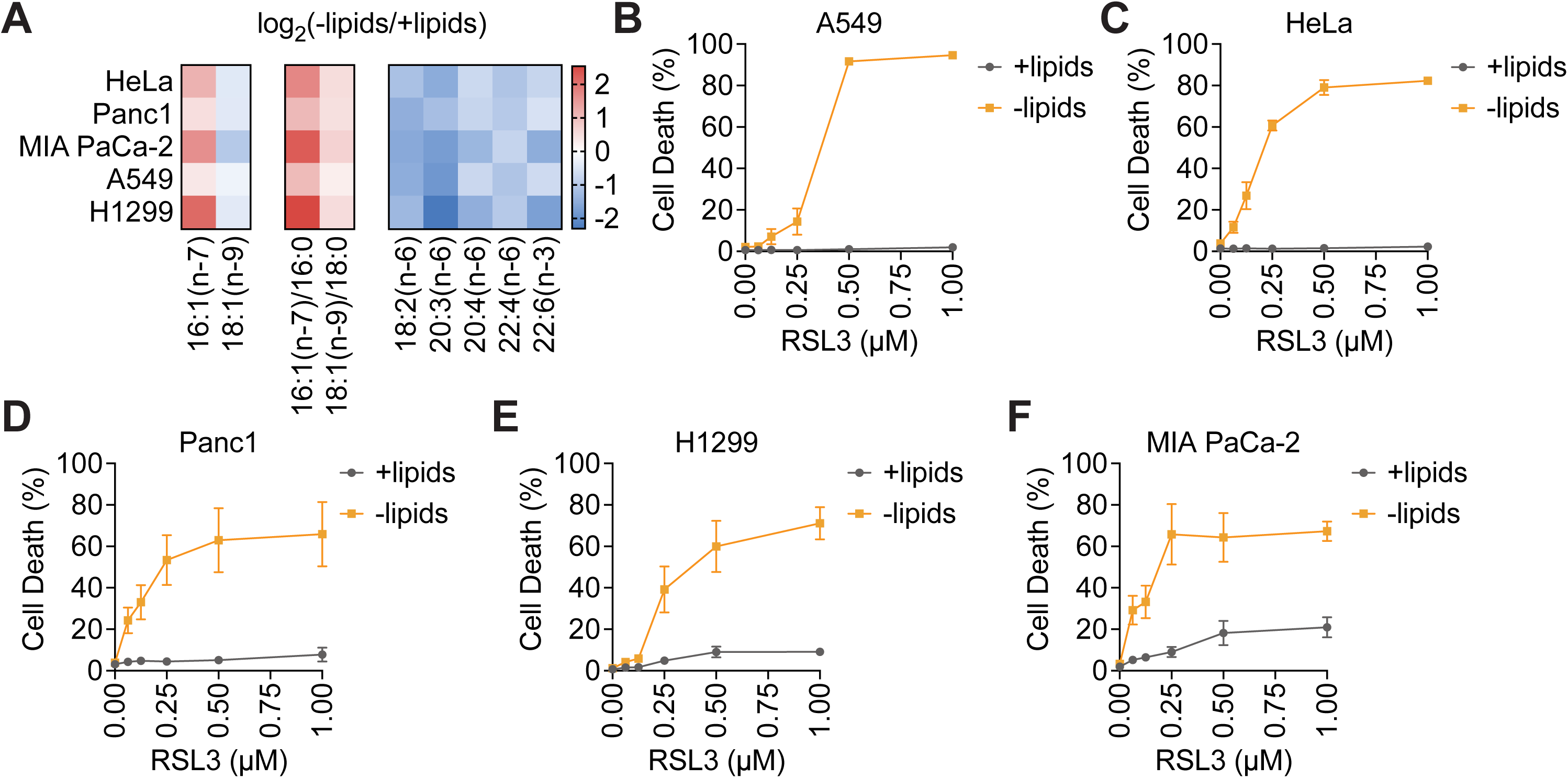
Lipid limitation sensitizes human cancer cells to RSL3-induced ferroptosis. **A,** Heat map showing fold changes in levels of the indicated fatty acids and fatty acid ratios in cells cultured in lipid-depleted versus lipid-replete media for 24 h. **B-F,** Cell death of A549 (**B**), HeLa (**C**), Panc1 (**D**), H1299 (**E**), and MIA PaCa-2 (**F**) cells treated with the indicated concentrations of RSL3 for 24 h after initial pre-treatment of cells in lipid-depleted versus lipid-replete media for 24 h. Data are presented as mean ± s.e.m; *n* = 3 biologically independent replicates.

High MUFA levels protect against ferroptosis by displacing PUFA-containing phospholipids from the plasma membrane^10^, whereas the PUFAs 20:4(n-6) and 22:4(n-6) are proposed to be the key PUFAs that are oxidized to trigger ferroptosis^5,6^. Because lipid-starved cancer cells have higher levels of 16:1(n-7) and lower levels of 20:4(n-6) and 22:4(n-6), we hypothesized that extracellular lipid limitation would cause cancer cells to be more resistant to ferroptosis. To test this, we treated our panel of cancer cell lines with the ferroptosis inducer RSL3, which is a covalent inhibitor of the lipid peroxide detoxifying enzyme GPX4. Surprisingly, lipid-starved cancer cells were significantly more sensitive to RSL3-induced ferroptosis despite having lower levels of PUFAs that can be oxidized (Fig. 1B-F). These results suggest that total cellular levels of MUFAs and PUFAs do not necessarily correlate with ferroptosis sensitivity.

### Lipid limitation induces the trafficking of PUFAs from triglycerides into ether phospholipids

To reconcile lower levels of oxidizable PUFAs with increased ferroptosis sensitivity, we asked whether PUFAs might accumulate in specific complex lipid classes in lipid-starved cancer cells. Using liquid chromatography-mass spectrometry (LC-MS), we conducted lipidomics in A549 and HeLa cells grown in lipid-replete versus lipid-depleted culture media. By first examining effects on total levels of each complex lipid class, we found that not all lipid classes changed in response to extracellular lipid limitation. For example, lipid-starved cells had reduced levels of storage lipids, such as triglycerides (TG) and cholesteryl esters (ChE), whereas levels of phospholipids, including phosphatidylethanolamine (PE) and phosphatidylcholine (PC), were maintained (Fig. 2A, S1A). This observation may be consistent with lipid-starved cells liberating fatty acids from storage lipids in order to maintain phospholipid levels.

**Figure 2.**
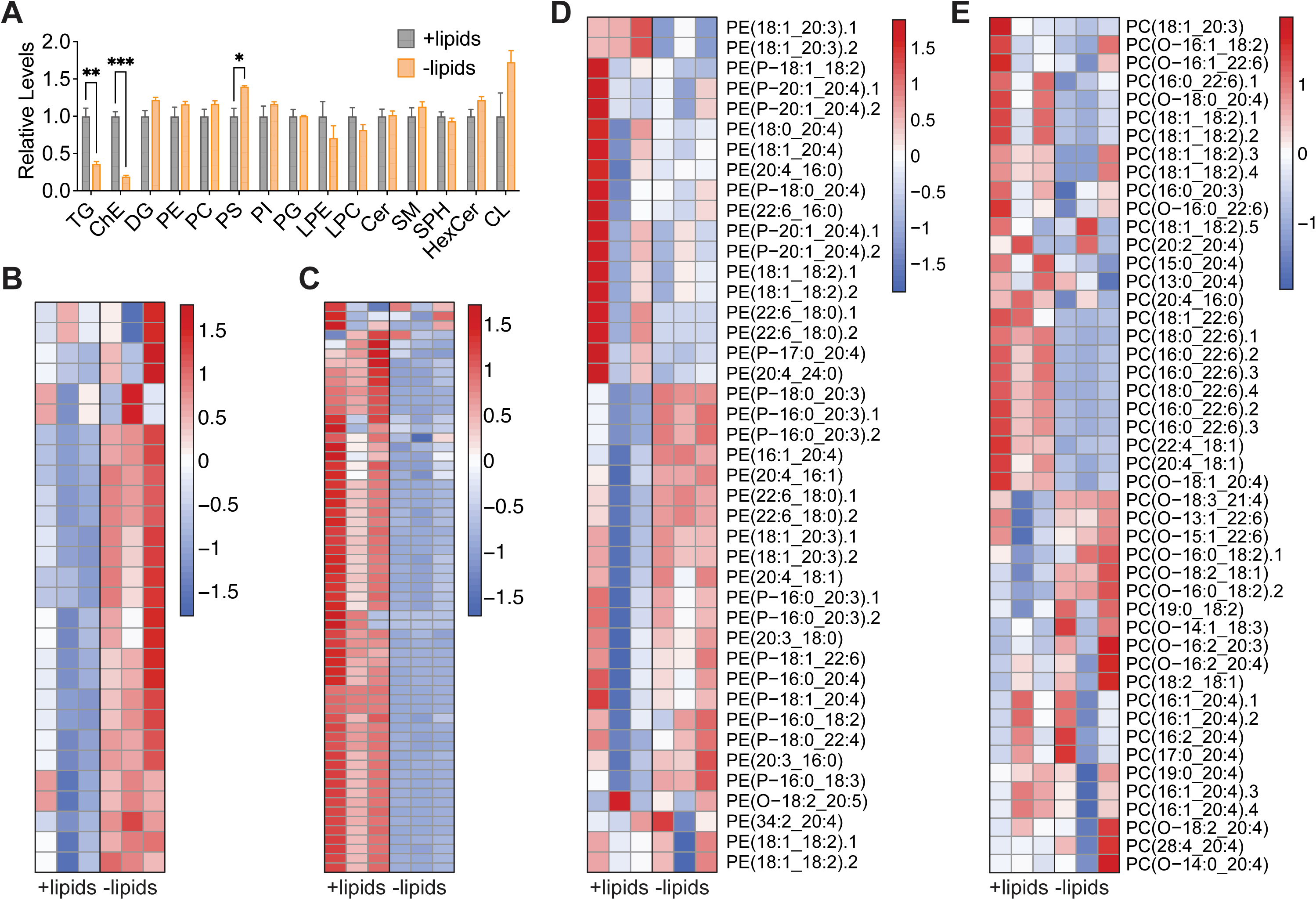
Lipid limitation decreases levels of storage lipids and increases levels of PUFA-containing ether PEs and PCs. A549 cells were cultured in lipid-depleted versus lipid-replete media for 24 h prior to extracting lipids for analysis by LC-MS. **A,** Relative levels of complex lipid classes. **B,** Heat map showing relative levels of phospholipids containing two MUFA side chains. **C,** Heat map showing relative levels of TGs containing at least one PUFA. **D,** Heat map showing relative levels of PEs containing at least one PUFA. **E,** Heat map showing relative levels of PCs containing at least one PUFA. TG, triglyceride; ChE, cholesterol ester; DG, diglyceride; PE, phosphatidylethanolamine; PC, phosphatidylcholine; PS, phosphatidylserine; PI; phosphatidylinositol; PG, phosphatidylglycerol; LPE, lysophosphatidylethanolamine; LPC, lysophosphatidylcholine; Cer, ceramide; SM, sphingomyelin; SPH, sphingosine; HexCer, hexosylceramide; CL, cardiolipin. Data are presented as mean ± s.e.m; *n* = 3 biologically independent replicates. Comparisons were made using a two-tailed Student’s t-test (**A**). \**P<0.05*, \*\**P<0.01*, \*\*\**P<0.001*.

We next examined how extracellular lipid limitation alters the distribution of MUFAs and PUFAs across different lipid classes. Consistent with increased MUFA levels and higher SCD activity in lipid-starved cells (Fig. 1A), levels of all phospholipids containing only MUFA chains were elevated in A549 and HeLa cells grown in lipid-depleted media (Fig. 2B, S1B). For PUFAs, while levels of PUFA-containing TGs were reduced in lipid-starved cells (Fig. 2C, Fig. S1C), PUFA-containing PEs and PCs exhibited more heterogeneity (Fig. 2D-E, S1D-E). Notably, the PUFA-containing PEs and PCs that were elevated in lipid-starved cells were enriched with 1) longer chain, highly unsaturated fatty acids such as 20:4 and 22:4, and 2) phospholipids with ether (O-) or vinyl ether (P-) acyl linkages. Taken together, these data suggest that in response to extracellular lipid limitation, cancer cells may initiate lipolysis to liberate PUFAs from TGs, metabolize these PUFAs to produce long chain, highly unsaturated fatty acids such as 20:4(n-6) and 22:4(n-6), and channel these PUFAs into the synthesis of ether PEs and PCs. Importantly, because 20:4(n-6) and 22:4(n-6) are proposed to be the key PUFAs that become oxidized to trigger ferroptosis^5,6^, and ether phospholipid synthesis was recently identified to be essential for ferroptosis in some cell contexts^14–16^, these results propose that lipid-starved cancer cells, despite having lower PUFA levels, are more sensitive to ferroptosis because they accumulate key PUFAs in ether phospholipids.

To test whether PUFAs are being trafficked through this route, we conducted a pulse-chase stable isotope fatty acid labeling experiment. Cancer cells were first exposed to uniformly labeled ^13^C-18:2(n-6) for 24 hours. The tracer was then washed off, and cells were grown in unlabeled lipid-replete versus lipid-depleted media for another 24 hours prior to lipid extraction, allowing us to evaluate how lipid availability influences the re-distribution of ^13^C-18:2(n-6) across the lipidome (Fig. 3A). For lipid extraction, we first used a three-phase liquid extraction protocol that allowed us to analyze a non-polar lipid fraction, which contains TGs, versus a polar lipid fraction that includes phospholipids^17^. GC-MS was then used to detect [M+18] labeled PUFAs in these fractions. In A549 cells, we found that during the pulse period, 18:2(n-6) and its downstream PUFAs each became labeled in both the non-polar and polar lipid fractions (Fig. 3B, C). During the chase period into lipid-replete versus lipid-depleted media, [M+18] PUFA levels in the non-polar fraction were reduced, suggesting that lipolysis and TG turnover occurred under both conditions (Fig. 3B). However, the disappearance of [M+18] PUFAs from the non-polar fraction was greater in lipid-starved cells (Fig. 3B), consistent with our hypothesis that extracellular lipid limitation stimulates TG lipolysis to liberate PUFAs. In the polar lipid fraction containing phospholipids, [M+18] PUFA levels dropped during the chase period in lipid-replete media (Fig. 3C). However, in lipid-starved cells, the levels of long chain, highly unsaturated [M+18] 20:4(n-6) and 22:4(n-6) accumulated in phospholipids, whereas the precursor [M+18] 18:2(n-6) was reduced (Fig. 3C). These results indicate that during lipid limitation, the PUFAs liberated from TGs are further elongated and desaturated to form long chain, highly unsaturated PUFAs that then accumulate in phospholipids. To confirm that these effects are not specific to A549 cells, we used the same tracing strategy to show increased TG lipolysis and 20:4(n-6) and 22:4(n-6) accumulation in phospholipids in HeLa (Fig. S2A, B) and Panc1 (Fig. S2C, D) cells.

**Figure 3.**
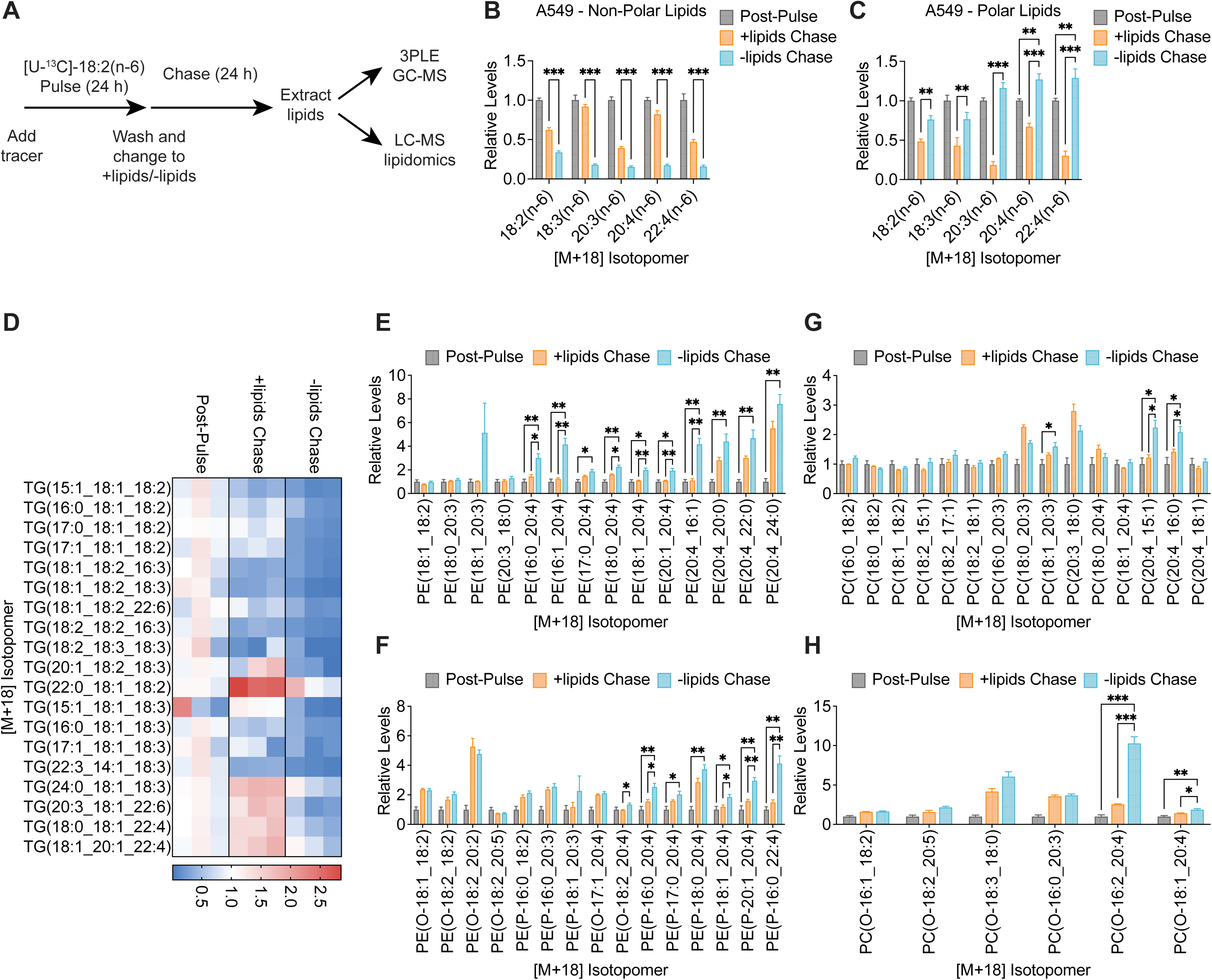
Lipid limitation induces the trafficking of PUFAs from triglycerides into ether phospholipids. **A,** Schematic illustrating pulse labeling of cells with [U-^13^C]-18:2(n-6), followed by a chase period into lipid-depleted versus lipid-replete media prior to lipid extraction and analysis. 3PLE, 3-phase liquid extraction; GC-MS, gas chromatography mass spectrometry; LC-MS, liquid chromatography mass spectrometry. **B, C,** Relative levels of [M+18] labeled PUFAs in the non-polar (**B**) or polar (**C**) lipid fractions from A549 cells post-pulse and after the chase period into lipid-depleted versus lipid-replete media. **D-H,** Relative levels of [M+18] labeled TGs (**D**), PEs (**E**), ether PEs (**F**), PCs (**G**), and ether PCs (**H**) containing at least one PUFA from A549 cells post-pulse and after the chase period into lipid-depleted versus lipid-replete media. Data are presented as mean ± s.e.m; *n* = 3 biologically independent replicates. Comparisons were made using two-way ANOVA (**B, C**) or a two-tailed Student’s t-test (**E-H**). \**P<0.05*, \*\**P<0.01*, \*\*\**P<0.001*.

Finally, we conducted the same pulse-chase experiment in A549 cells and analyzed lipid extracts by LC-MS to determine [M+18] PUFA labeling in complex lipids (Fig. 3A). During the chase period, levels of [M+18] TGs containing PUFAs decreased to a greater extent in cells growing in lipid-depleted versus lipid-replete media, confirming increased TG lipolysis in lipid-starved cells (Fig. 3D). In contrast, levels of [M+18] PEs, PCs, ether PEs, and ether PCs containing specifically 20:4(n-6) and 22:4(n-6) accumulated more in lipid-starved cells (Fig. 3E-H). Taken together, these results support our model that extracellular lipid limitation stimulates the liberation of PUFAs from TGs, which are then elongated and desaturated into long chain, highly unsaturated PUFAs that are used to synthesize phospholipids, including ether phospholipids. Because oxidative damage of PUFA-containing phospholipids specifically triggers ferroptosis^1,2^, we propose that lipid-starved cancer cells exhibit increased sensitivity to ferroptosis despite having overall lower PUFA levels because they accumulate PUFAs in their phospholipid pool.

### Triglyceride lipolysis is required for ferroptosis sensitivity in lipid-starved cancer cells

To test whether this trafficking of PUFAs in lipid-starved cells contributes to increased ferroptosis sensitivity, we first asked whether TG lipolysis is required for this effect. TG lipolysis is initiated by adipose triglyceride lipase (ATGL), which catalyzes the rate-limiting hydrolysis of TGs to generate diglycerides (DGs) and a non-esterified fatty acid. Hormone-sensitive lipase (HSL) and monoglyceride lipase (MGL) then hydrolyze DGs into monoglycerides (MGs), and MGs into glycerol, respectively^18^ (Fig. 4A). Because TG depletion occurs in lipid-starved cells (Fig. 2A), we first measured *ATGL* gene expression and found no change across all five cell lines in response to extracellular lipid limitation (Fig. 4B). We also found that ATGL protein levels decreased in lipid-starved A549 and HeLa cells and did not change in lipid-starved Panc1 cells (Fig. 4C). These results suggest that lipid-starved cells do not increase TG lipolysis by upregulating ATGL transcript or protein levels.

**Figure 4.**
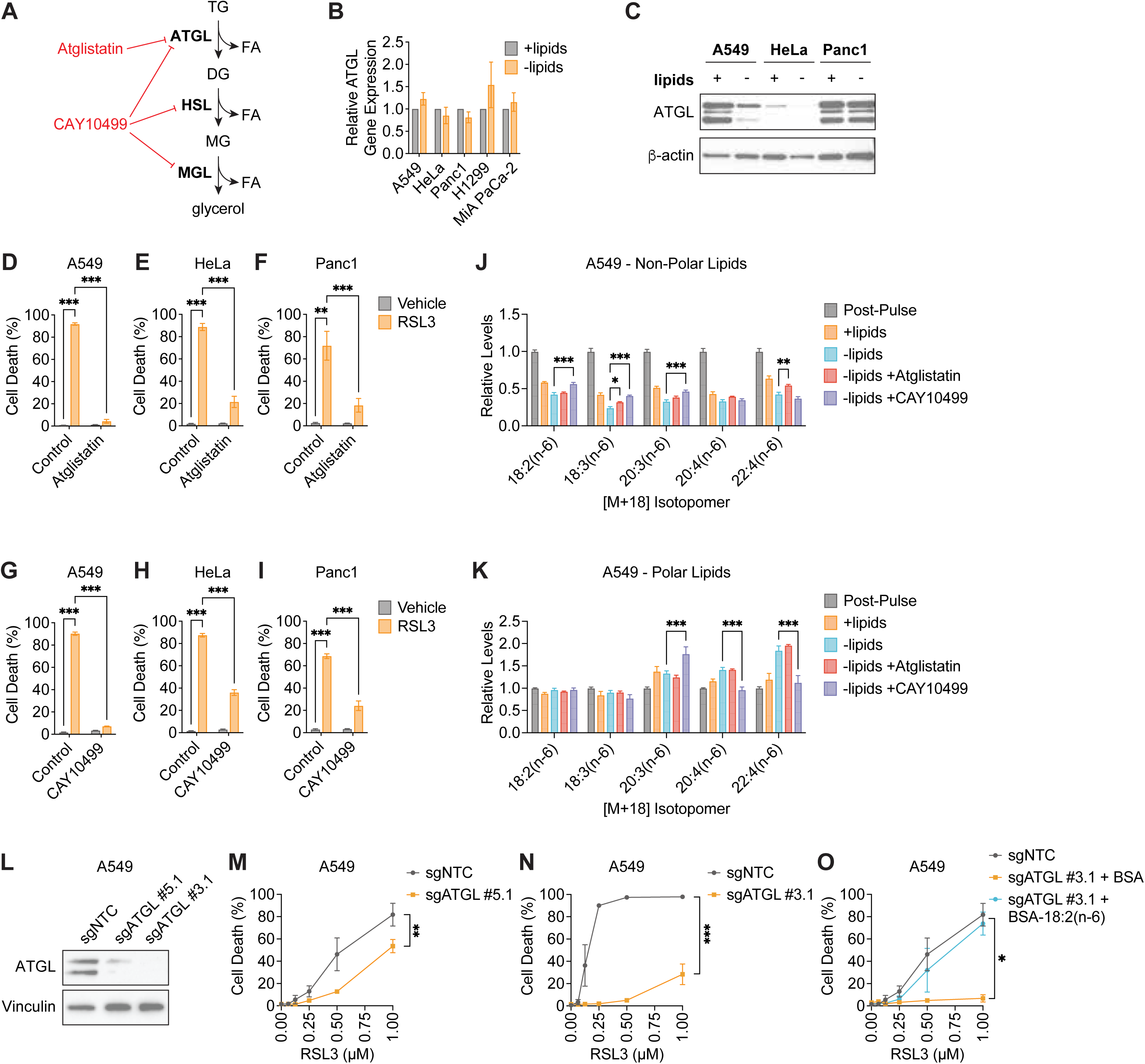
Inhibiting triglyceride lipolysis suppresses ferroptosis in lipid-starved cancer cells. **A,** Schematic illustrating the TG lipolysis pathway and lipase inhibitors. TG, triglyceride; DG, diglyceride; MG, monoglyceride; FA, fatty acid; ATGL, adipose triaglyceride lipase; HSL, hormone sensitive lipase; MGL, monoglyceride lipase. **B,** Relative *ATGL* gene expression in the indicated cells cultured in lipid-depleted versus lipid-replete media for 24 h. **C,** Immunoblot for ATGL and β-actin in the indicated cells cultured in lipid-depleted versus lipid-replete media for 24 h. **D-F,** Cell death of A549 (**D**), HeLa (**E**), and Panc1 (**F**) cells treated with 1 µM RSL3 for 24 h after initial pre-treatment of cells in lipid-depleted versus lipid-replete media, with or without 10 µM Atglistatin, for 24 h. **G-I,** Cell death of A549 (**G**), HeLa (**H**), and Panc1 (**I**) cells treated with 1 µM RSL3 for 24 h after initial pre-treatment of cells in lipid-depleted versus lipid-replete media, with or without 5 µM CAY10499, for 24 h. **J, K,** Cells were pulse labeled with [U-^13^C]-18:2(n-6), followed by a chase period prior to lipid extraction and analysis, as shown in Fig. 3A. Relative levels of [M+18] labeled PUFAs in the non-polar (**J**) or polar (**K**) lipid fractions from A549 cells post-pulse and after the chase period into lipid-replete media or lipid-depleted media, with or without 10 µM Atglistatin or 5 µM CAY10499. **L,** Immunoblot for ATGL and vinculin in A549 cells in which ATGL was knocked out by CRISPR-Cas9 with two independent sgRNAs. **M, N,** Cell death of A549 sgNTC cells versus sgATGL #5.1 (**M**) or sgATGL #3.1 (**N**) cells treated with the indicated concentrations of RSL3 for 24 h after initial pre-treatment of cells in lipid-depleted media for 24 h. **O,** Cell death of A549 sgNTC versus sgATGL #3.1 cells treated with the indicated concentrations of RSL3 for 24 h after initial pre-treatment of cells in lipid-depleted media containing bovine serum albumin (BSA) control or 25 µM BSA-conjugated 18:2(n-6) for 24 h. Data are presented as mean ± s.e.m; *n* = 3 biologically independent replicates. Comparisons were made using two-way ANOVA (**D-K, M-O**). \**P<0.05*, \*\**P<0.01*, \*\*\**P<0.001*.

To test if TG lipolysis is necessary for ferroptosis sensitivity, we treated lipid-starved cancer cells with Atglistatin^19^, an ATGL inhibitor, and CAY10499^20^, a non-selective lipase inhibitor that targets ATGL, HSL, and MGL (Fig. 4A). We found that in A549, HeLa, and Panc1 cells, RSL3-induced ferroptosis was prevented by both Atglistatin (Fig. 4D-F) and CAY10499 (Fig. 4G-I). To determine whether this was due to on-target inhibition of TG lipolysis by Atglistatin and CAY10499, we conducted the pulse-chase ^13^C-18:2(n-6) labeling experiment described in Fig. 3A to determine how Atglistatin and CAY10499 alter the re-distribution of [M+18] PUFAs in the non-polar versus polar lipid fractions of lipid-starved cells. Atglistatin minimally prevented the depletion of [M+18] PUFAs from the non-polar lipid fraction in lipid-starved A549, HeLa, and Panc1 cells, and also did not block the accumulation of [M+18] 20:4(n-6) and 22:4(n-6) in the polar lipid fraction (Fig. 4J-K, Fig. S3A-D). This suggests that Atglistatin is not an effective ATGL inhibitor in these human cell lines, as has been suggested in previous studies^20,21^, and may be preventing ferroptosis through another mechanism. In contrast, CAY10499 prevented the depletion of [M+18] PUFAs, particularly 18:2(n-6), in the non-polar lipid fraction from lipid-starved cells, and also blocked the accumulation of [M+18] 20:4(n-6) and 22:4(n-6) in the polar lipid fraction (Fig. 4J-K, Fig. S3A-D). Therefore, CAY10499 impaired TG lipolysis and prevented ferroptosis in lipid-starved cells.

To further validate these findings, we used CRISPR-Cas9 to knock out *ATGL* in A549 cells with two distinct single guide RNAs (sgRNAs) (Fig. 4L). Consistent with pharmacological inhibition of TG lipolysis, we found that loss of *ATGL* in A549 cells promoted resistance to RSL3- induced ferroptosis under lipid-limited conditions (Fig. 4M-N). Finally, based on our model, *ATGL* loss should prevent the liberation of PUFAs from TGs in lipid-starved cells. Therefore, we asked whether supplementing *ATGL* knockout cells with 18:2(n-6), the precursor to 20:4(n-6) and 22:4(n-6), could restore ferroptosis sensitivity. Indeed, A549 *ATGL* knockout cells were susceptible to RSL3-induced ferroptosis when 18:2(n-6) was supplied in lipid-depleted culture media (Fig. 4O). Together, these data are consistent with our model that TG lipolysis is required to sensitize cancer cells to ferroptosis under lipid-limited conditions.

### Increased PUFA metabolism is necessary for ferroptosis sensitivity in lipid-starved cancer cells

We next asked whether the production of long chain, highly unsaturated PUFAs such as 20:4(n-6) and 22:4(n-6) is needed for increased ferroptosis sensitivity in lipid-starved cells. Though mammalian cells cannot synthesize PUFAs *de novo*, they can elongate and further desaturate PUFAs taken up from the environment, such as 18:2(n-6), through the fatty acid elongases ELOVL2/5 and desaturases FADS1/2 (Fig. 5A). To evaluate how extracellular lipid limitation affects the PUFA metabolism pathway, we first measured the gene expression of *ELOVL2*, *ELOVL5*, *FADS1*, and *FADS2* in A549, HeLa, Panc1, H1299, and MIA PaCa-2 cells.

**Figure 5.**
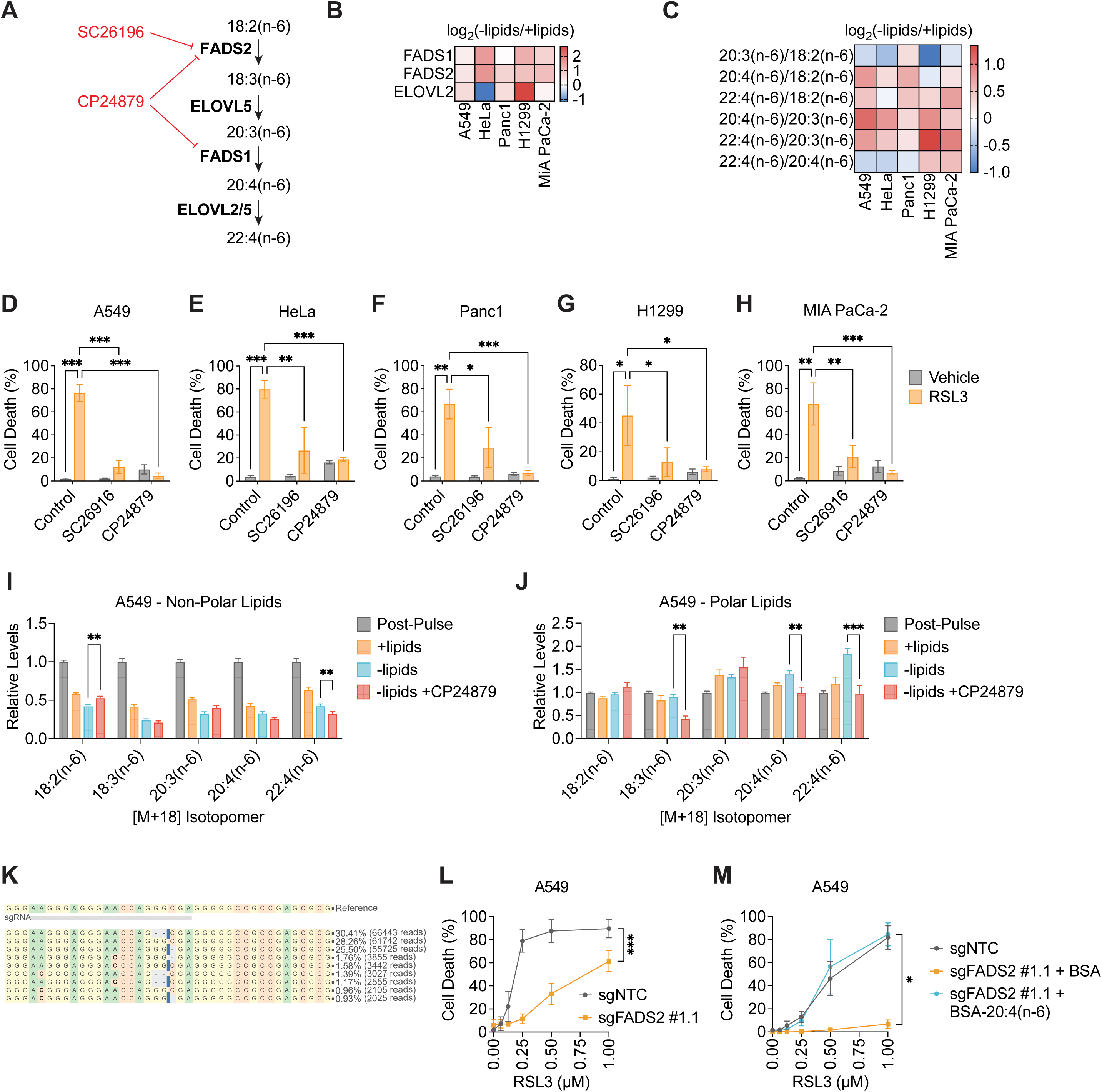
Inhibiting PUFA metabolism suppresses ferroptosis in lipid-starved cancer cells. **A,** Schematic illustrating the PUFA metabolism pathway and inhibitors of FADS1 and FADS2. FADS2, fatty acid desaturase 2; ELOVL2/5, elongase of very-long fatty acid 2/5; FADS1, fatty acid desaturase 1. **B,** Fold change in gene expression of FADS1, FADS2, and ELOVL2 in the indicated cells cultured in lipid-depleted versus lipid-replete media for 24 h. **C,** Fold change in the relative ratios of highly unsaturated PUFAs to their precursor PUFAs in the indicated cells cultured in lipid-depleted versus lipid-replete media for 24 h. **D-H,** Cell death of A549 (**D**), HeLa (**E**), Panc1 (**F**), H1299 (**G**), and MIA PaCa-2 (**H**) cells treated with 1 µM RSL3 for 24 h after initial pre-treatment of cells in lipid-depleted versus lipid-replete media, with or without 5 µM SC26196 or 10 µM CP24879, for 24 h. **I, J,** Cells were pulse labeled with [U-^13^C]-18:2(n-6), followed by a chase period prior to lipid extraction and analysis, as shown in Fig. 3A. Relative levels of [M+18] labeled PUFAs in the non-polar (**I**) or polar (**J**) lipid fractions from A549 cells post-pulse and after the chase period into lipid-replete media or lipid-depleted media, with or without 10 µM CP24879. **K,** Sequencing confirmation of *FADS2* gene editing by CRISPR-Cas9 in A549 cells. Indel sequences and allele frequencies taken from analysis using CRISPResso2^40^. **L,** Cell death of A549 sgNTC cells versus sgFADS2 #1.1 cells treated with the indicated concentrations of RSL3 for 24 h after initial pre-treatment of cells in lipid-depleted media for 24 h. **M,** Cell death of A549 sgNTC versus sgFADS2 #1.1 cells treated with the indicated concentrations of RSL3 for 24 h after initial pre-treatment of cells in lipid-depleted media containing bovine serum albumin (BSA) control or 25 µM BSA-conjugated 20:4(n-6) for 24 h. Data are presented as mean ± s.e.m; *n* = 3 biologically independent replicates. Comparisons were made using two-way ANOVA (**D-J, L-N**). \**P<0.05*, \*\**P<0.01*, \*\*\**P<0.001*.

While *ELOVL5* expression was undetectable in these cells, we found that lipid starvation increased expression of *FADS1*, *FADS2*, and *ELOVL2* in all five cell lines (Fig. 5B). We also examined how extracellular lipid limitation alters the relative levels of long chain, highly unsaturated PUFAs in cancer cells. As described earlier, lipid-starved cancer cells have overall lower levels of PUFAs (Fig. 1A). However, the ratios of the levels of each PUFA to the levels of its upstream PUFAs (e.g. the 20:4(n-6)/18:2(n-6) ratio) can be reflective of PUFA metabolism activity. Interestingly, we find that many of these PUFA ratios were elevated by extracellular lipid limitation across all five cancer cell lines (Fig. 5C). These results suggest that even though lipid-starved cancer cells have lower total PUFA levels, a higher proportion of the PUFAs present within these cells are long chain, highly unsaturated fatty acids such as 20:4(n-6) and 22:4(n-6). These observations are consistent with data from our pulse-chase ^13^C-18:2(n-6) labeling experiment. In lipid-starved cancer cells, levels of [M+18] 18:2(n-6) and 18:3(n-6) dropped in the phospholipid fraction during the chase period, whereas levels of [M+18] 20:4(n-6) and 22:4(n-6) rose (Fig. 3C), which likely resulted from increased elongation and desaturation of the supplied ^13^C-18:2(n-6). Taken together, these data suggest that extracellular lipid limitation stimulates PUFA metabolism pathway activity.

Because highly unsaturated PUFAs are key fatty acid species that become oxidized to trigger ferroptosis^5,6^, we asked whether up-regulated PUFA metabolism is required for lipid limitation to increase ferroptosis sensitivity. We treated lipid-starved cancer cells with the FADS2 inhibitor SC26196^22,23^ or the dual FADS1/FADS2 inhibitor CP24879^23,24^ before inducing ferroptosis with RSL3. Across all five cell lines, we observed that both FADS2 inhibition and FADS1/2 dual inhibition prevented ferroptosis in lipid-starved cancer cells (Fig. 5D-H). We also conducted a pulse-chase ^13^C-18:2(n-6) labeling experiment to determine how SC26196 and CP24879 alter the re-distribution of [M+18] PUFAs in the non-polar versus polar lipid fractions of lipid-starved cells. In A549 cells, both SC26916 and CP24879 did not strongly prevent the reduction of [M+18] PUFAs from the TG fraction of lipid-starved cells, but did block the accumulation of [M+18] 20:4(n-6) and 22:4(n-6) in the phospholipid fraction (Fig. 5I-J, Fig. S4A-B). Similar results were observed for HeLa and Panc1 cells (Fig. S4C-J).

We also used CRISPR-Cas9 to knock out *FADS2* in A549 cells. We were unable to identify an appropriate antibody to screen for knockout clones by immunoblotting. However, we used next-generation sequencing to identify one A549 *FADS2* knockout clone in which the majority of *FADS2* alleles were modified by CRISPR-Cas9 editing of exon 1 (Fig. 5K). We found that similar to FADS2 inhibition, loss of *FADS2* promoted ferroptosis resistance under lipid-limited conditions (Fig. 5L). As the first step in the PUFA metabolism pathway, FADS2 is required for the production of 20:4(n-6) and 22:4(n-6) from 18:2(n-6) (Fig. 5A). Consistently, supplementing *FADS2* knockout cells with extracellular 20:4(n-6) in lipid-depleted media restored their sensitivity to ferroptosis (Fig. 5M). Taken together, these data support our model that increased PUFA metabolism pathway activity is needed for lipid limitation to stimulate 20:4(n-6) and 22:4(n-6) production for phospholipid synthesis, which subsequently contributes to increased ferroptosis sensitivity.

### Ether lipid synthesis is required for ferroptosis sensitivity in lipid-starved cancer cells

Finally, because levels of ether PEs and PCs containing 20:4(n-6) and 22:4(n-6) were elevated in lipid-starved cancer cells (Fig. 2D-E, Fig. 3G-H, Fig. S1D-E), we asked whether ether phospholipid synthesis contributes to increased ferroptosis sensitivity under extracellular lipid limitation. Ether lipid synthesis begins in the peroxisome, where fatty alcohols produced by fatty acyl-CoA reductase (FAR1) are attached to a dihydroxyacetone phosphate backbone by alkylglycerone phosphate synthase (AGPS) to form alkyl-DHAP. Alkyl-DHAP is then reduced to 1-O-alkyl-glycerol-3-phophate before being transferred to the endoplasmic reticulum (ER) for further processing (Fig. 6A). Across all five cancer cell lines, we found no change in *AGPS* and *FAR1* gene expression (Fig. 6B-C) or in AGPS protein levels (Fig. 6D) in lipid-starved cells. To test if ether lipid synthesis is necessary for increased ferroptosis sensitivity, we treated lipid-starved cancer cells with the AGPS inhibitor AGPS-IN-2i^25^ before inducing ferroptosis with RSL3. Indeed, AGPS-IN-2i prevented ferroptosis in lipid-starved A549, HeLa, and Panc1 cells (Fig. 6E-G).

**Figure 6.**
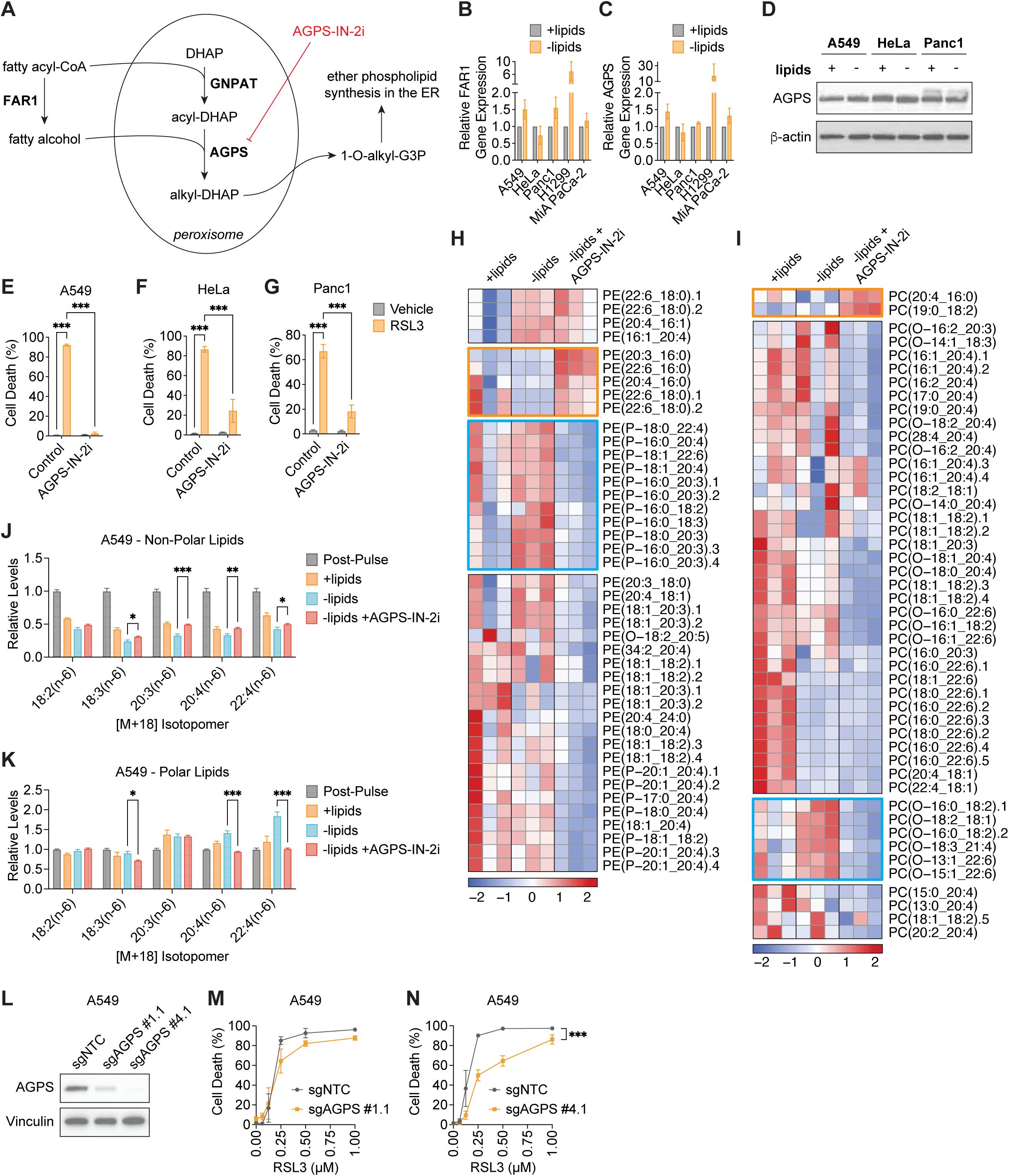
Inhibiting ether lipid synthesis suppresses ferroptosis in lipid-starved cells. **A,** Schematic illustrating the ether lipid synthesis pathway and an AGPS inhibitor. FAR1, fatty acyl-CoA reductase 1; GNPAT, glyceronephosphate O-acyltransferase; AGPS, alkylglycerone phosphate synthase; DHAP, dihydroxyacetone phosphate; G3P, glycerol 3-phosphate; ER, endoplasmic reticulum. **B, C,** Relative *FAR1* (**B**) and *AGPS* (**C**) gene expression in the indicated cells cultured in lipid-depleted versus lipid-replete media for 24 h. **D,** Immunoblot for AGPS and β-actin in the indicated cells cultured in lipid-depleted versus lipid-replete media for 24 h. **E-G,** Cell death of A549 (**E**), HeLa (**F**), and Panc1 (**G**) cells treated with 1 µM RSL3 for 24 h after initial pre-treatment of cells in lipid-depleted versus lipid-replete media, with or without 150 µM AGPS-IN-2i, for 24 h. **H, I,** Heat map showing relative levels of PEs (**H**) or PCs (**I**) containing at least one PUFA in A549 cells cultured in lipid-replete or lipid-depleted media, with or without 150 µM AGPS-IN-2i, for 24 h. Blue boxes highlight ether phospholipid species that decrease upon AGPS-IN-2i treatment. Orange boxes highlight diacyl phospholipid species that increase upon AGPS-IN-2i treatment. **J, K,** Cells were pulse labeled with [U-^13^C]-18:2(n-6), followed by a chase period prior to lipid extraction and analysis, as shown in Fig. 3A. Relative levels of [M+18] labeled PUFAs in the non-polar (**J**) or polar (**K**) lipid fractions from A549 cells post-pulse and after the chase period into lipid-replete media or lipid-depleted media, with or without 150 µM AGPS-IN-2i. **L,** Immunoblot for AGPS and vinculin in A549 cells in which AGPS was knocked out by CRISPR-Cas9 with two independent sgRNAs. **M, N,** Cell death of A549 sgNTC cells versus sgAGPS #1.1 (**M**) or sgAGPS #4.1 (**N**) cells treated with the indicated concentrations of RSL3 for 24 h after initial pre-treatment of cells in lipid-depleted media for 24 h. Data are presented as mean ± s.e.m; *n* = 3 biologically independent replicates. Comparisons were made using two-way ANOVA (**E-G, J, K, M, N**). \**P<0.05*, \*\**P<0.01*, \*\*\**P<0.001*.

To confirm the on-target effects of AGPS-IN-2i, we used LC-MS to evaluate how AGPS-IN-2i alters the ether phospholipid composition of lipid-starved cancer cells. We found that the accumulation of PUFA-containing ether PEs and ether PCs in lipid-starved cells was indeed reversed with AGPS-IN-2i treatment in A549 cells (Fig. 6H-I, blue boxes). Interestingly, AGPS-IN-2i increased the levels of some diacyl PE and PC species containing PUFAs in lipid-starved A549 cells (Fig. 6H-I, orange boxes), suggesting that preventing PUFA utilization for ether phospholipid synthesis may divert PUFAs towards diacyl phospholipid synthesis. Similar results were observed in lipid-starved HeLa cells treated with AGPS-IN-2i (Fig. S5A-B). These observations indicate that AGPS-IN-2i effectively prevented the accumulation of PUFA-containing ether phospholipids in lipid-starved cells.

We also conducted a pulse-chase ^13^C-18:2(n-6) labeling experiment to determine how AGPS-IN-2i alters the re-distribution of [M+18] PUFAs in the non-polar versus polar lipid fractions of lipid-starved cells. AGPS-IN-2i completely prevented the accumulation of [M+18] 20:4(n-6) and 22:4(n-6) in the phospholipid fraction of lipid-starved A549 cells (Fig. 6J-K), suggesting that a significant proportion of [M+18] PUFAs that accumulate in phospholipids in lipid-starved cells were contained in ether phospholipids. Similar results were observed in HeLa and Panc1 cells (Fig. S5C-F). Finally, we knocked out *AGPS* in A549 cells using CRISPR-Cas9 with two distinct sgRNAs (Fig. 6L). A549 sgAGPS #1.1 cells retained some residual AGPS expression and were only slightly more resistant to RSL3 (Fig. 6M), whereas A549 sgAGPS #4.1 cells showed complete knockout and were more resistant to ferroptosis (Fig. 6N). Together, these data are consistent with our model that the channeling of long chain, highly unsaturated PUFAs into ether phospholipids in cancer cells contributes to their increased sensitivity to ferroptosis under lipid-limited conditions.

## DISCUSSION

Our data proposes a model for how extracellular lipid availability influences cancer cell sensitivity to ferroptosis (Fig. 7). When fatty acids are available to be taken up from the environment, they can be used for the formation of complex lipids, membrane synthesis, or bioenergetics, and excess fatty acids may be stored in complex lipids such as TGs. Under lipid-limited conditions, cancer cells can still obtain sufficient SFAs and MUFAs though *de novo* lipogenesis. However, because mammalian cells cannot *de novo* synthesize PUFAs, lipid-starved cancer cells increase TG lipolysis to liberate stored PUFAs. Moreover, PUFA metabolism is upregulated, such that non-esterified PUFAs are desaturated and elongated to generate long chain, highly unsaturated PUFAs such as 20:4(n-6) and 22:4(n-6). These PUFAs are then incorporated into phospholipids, particularly PE and PC ether lipids, which promote increased sensitivity to ferroptosis. These results illustrate that the total cellular levels of PUFAs versus MUFAs alone do not necessarily correlate with ferroptosis susceptibility in cancer cells. Rather, the intracellular trafficking of highly unsaturated PUFAs into the proper phospholipid pools is needed for PUFAs to promote ferroptosis.

**Figure 7.**
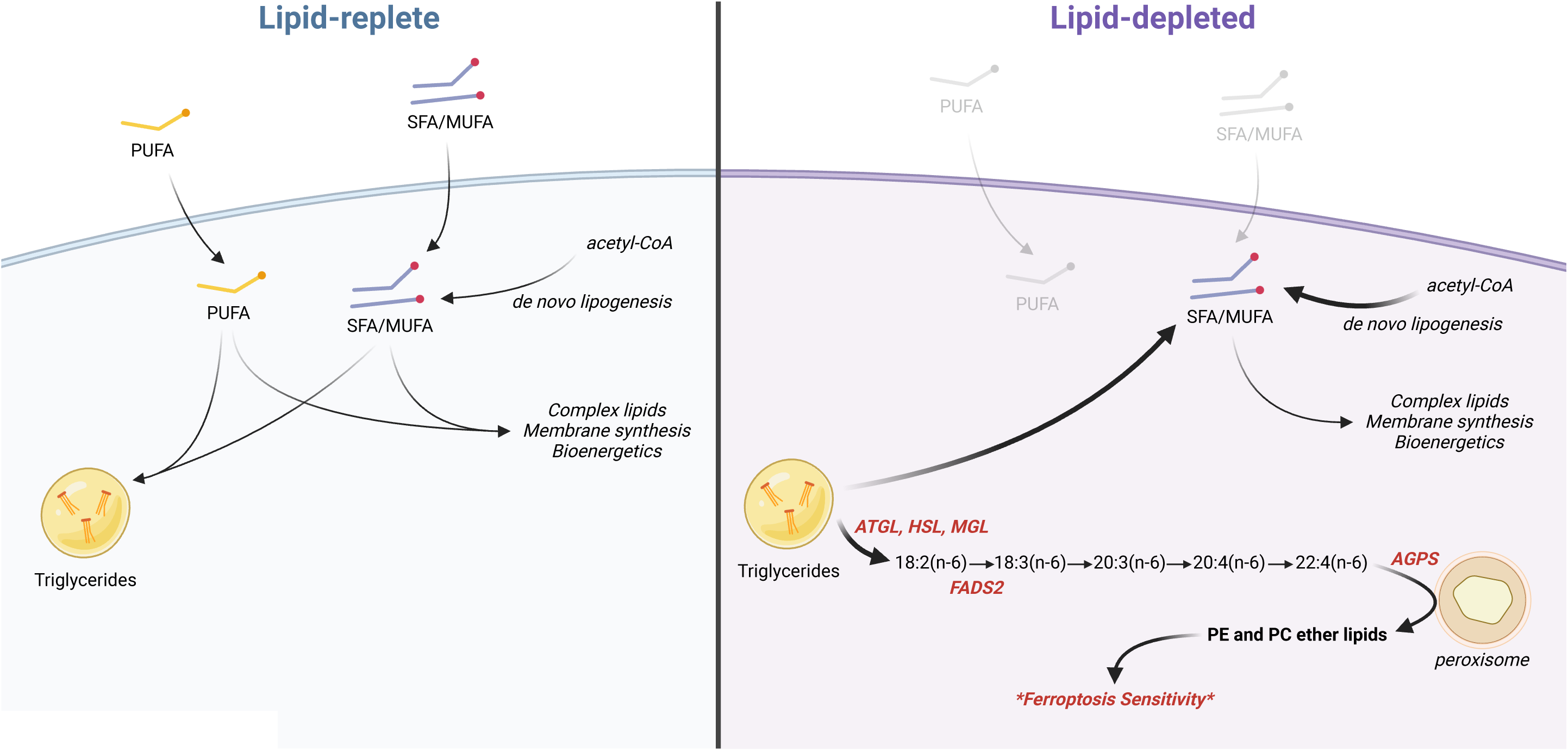
Proposed model for how extracellular lipid limitation increases ferroptosis susceptibility in cancer cells. Under lipid-replete conditions (left), fatty acids can be taken up from the environment to support cellular processes such as complex lipid synthesis, membrane synthesis, and bioenergetics. Excess fatty acids may be stored in storage lipids such as triglycerides. Under lipid-depleted conditions (right), PUFAs are liberated from triglycerides and further elongated and desaturated to generate highly unsaturated, long chain PUFAs such as 20:4(n-6) and 22:4(n-6). These PUFAs are then used to synthesize phospholipids, particularly ether PEs and PCs. Increased PUFA accumulation in the ether phospholipid pool results in increased ferroptosis sensitivity.

The mechanisms by which cancer cells sense reduced lipid availability to promote this intracellular PUFA trafficking pathway remain to be determined. We show that the stimulation of TG lipolysis does not involve induction of the transcript or protein levels of the rate-limiting lipolytic gene *ATGL* in lipid-starved cells (Fig. 4B-C). ATGL activity can also be altered by post-translational modifications and binding to co-regulatory factors^18^, and it is possible that these other regulatory mechanisms may be sensitive to extracellular lipid availability. For increased PUFA metabolism, we show that the PUFA desaturases FADS1 and FADS2 are upregulated at the transcript level across all cell lines analyzed (Fig. 5B). This likely occurs through lipid limitation inducing the activity of the SREBP transcription factors, which target both *FADS1* and *FADS2*^26^. Finally, how lipid-starved cells increase the incorporation of highly unsaturated PUFAs into ether phospholipids remains unclear, and likely does not involve transcriptional upregulation or increased protein levels of the ether lipid synthesis genes *AGPS* and *FAR1* (Fig. 6B-D).

Our results reinforce the growing recognition that the regulators of ferroptosis sensitivity are context dependent. A recent study that conducted an integrated analysis of 24 published genetic loss-of-function screens for ferroptosis regulators found little overlapping hits between the screens, suggesting that genes dictating ferroptosis sensitivity may vary between different cell lines and conditions^16^. Indeed, *ATGL* and *FADS1/2* were not identified as ferroptosis suppressors in these screens. However, these genes have been implicated in ferroptosis sensitivity in specific contexts: *ATGL* knockdown by siRNA was reported to suppress the toxicity of the GPX4 inhibitor ML162 in two human breast cancer cell lines^27^, and inhibition of *FADS2* or genetic knockout of *FADS1* in mesenchymal-type gastric cancer cells suppressed the toxicity of RSL3^28^. Our data elucidates the mechanisms by which TG lipolysis and the PUFA metabolism pathway participates in a PUFA trafficking pathway to promote ferroptosis sensitivity under lipid-limited conditions. Similarly, while some screens have identified *AGPS* as a ferroptosis suppressor gene^14–16^, AGPS-dependent ether lipid synthesis was shown to be dispensable for ferroptosis in some contexts, such as in HAP1 and HT-1080 cells^16^. It was shown that *AGPS* knockout re-routed PUFAs from ether lipids into diacyl phospholipids, and elevated levels of PUFA-containing diacyl phospholipids were sufficient in these cells to initiate ferroptosis^16^. In contrast, while our results also show the same diversion of PUFAs into diacyl phospholipids upon AGPS inhibition (Fig. 6H-I, Fig. S5A-B), AGPS is required for enhanced ferroptosis sensitivity in lipid-starved cancer cells, suggesting that higher levels of PUFA-containing diacyl phospholipids are not always sufficient to initiate ferroptosis.

Ferroptosis induction is being actively explored as a potential therapeutic strategy for targeting cancer cells. Understanding the mechanisms that regulate ferroptosis sensitivity under different conditions will be important for identifying contexts under which ferroptosis inducers will be effective, particularly in light of recent efforts to define and manipulate metabolite levels in the tumor microenvironment^29,30^. Our results add to growing evidence that manipulating extracellular lipid availability may be a strategy for enhancing cancer cell ferroptosis susceptibility. We recently showed that a calorie-restricted diet can reduce lipid availability in the tumor microenvironment^11^, and it is possible that such a diet may improve the efficacy of ferroptosis inducers. Similarly, high fat diets may also alter tumor ferroptosis sensitivity depending on the types of fatty acids that become elevated in the tumor microenvironment. For example, because *in vitro* supplementation of PUFAs, such as arachidonic acid, increases cancer cell susceptibility to ferroptosis, it has been shown that diets high in PUFAs can lead to ferroptosis induction in mouse tumor models^31^. However, because our results highlight that environmental and intracellular fatty acid levels alone do not necessarily correlate with ferroptosis susceptibility, it will be important to evaluate how different dietary interventions with distinct fatty acid compositions alter the trafficking and distribution of MUFAs and PUFAs across TGs, phospholipids, and ether phospholipids. Finally, many of these ideas remain difficult to test in mouse models of cancer because most existing ferroptosis inducers are unsuitable for *in vivo* use due to poor bioavailability. However, candidate compounds with better bioavailability are beginning to emerge^32–34^, and defining how dietary manipulation of fatty acid and lipid levels in the tumor microenvironment influences the efficacies of these drugs may facilitate better use of ferroptosis inducers for cancer treatment.

## Acknowledgements

We thank members of the Lien laboratory and Matthew Vander Heiden’s laboratory for useful discussions and experimental advice. E.C.L was supported by the Van Andel Institute MeNu Program and the NIH (R00CA255928).

## Author contributions

E.C.L. conceived the project. K.H.S., C.J.L., A.W., S.M., S.R.D., X.Y., M.O., and B.R.F. performed the experiments and analyzed data. K.H.S., T.J.R., and A.W. assisted with generating gene knockout cell lines. A.E.E., A.J., and R.D.S. assisted with mass spectrometry analyses of fatty acids and lipids. K.H.S. and E.C.L. wrote the manuscript with input from all authors.

## MATERIALS AND METHODS

### Cell lines and culture

All cell lines for this study were obtained from the American Type Culture Association. No cell lines used in this study were found in the database of commonly misidentified cell lines that is maintained by the International Cell Line Authentication Committee. Cells were regularly assayed for mycoplasma contamination and passaged for no more than 6 months. All cells were cultured in DMEM (Corning Life Sciences, 10-013-CV) without pyruvate and 10% heat-inactivated dialyzed fetal bovine serum (FBS) (Corning Life Sciences, 35-010-CV) unless otherwise specified. For lipid limitation experiments, FBS was stripped of lipids and dialyzed as previously described to generate lipid-depleted cell culture media^13^.

### Inhibitors

RSL3 (MilliporeSigma, SML2234), Atglistatin (Cayman Chemical, 15284), CAY10499 (Cayman Chemical, 10007875), SC26196 (Tocris, 4189), CP24879 (MilliporeSigma, C9115), and AGPS-IN-2i (Glixx Laboratories, GLXC-06522) were used at the indicated concentrations.

### Cell Death Assays

Cells were seeded at an initial density of 20,000-50,000 cells per well on a 24-well plate in 1 ml of DMEM medium. After incubating for 24 hours, cells were washed three times with 1 ml of PBS and changed to the indicated growth conditions and drug treatments for 24 hours. Cells were then treated with the indicated concentrations of RSL3 and 20 nM Sytox Green (Invitrogen, S7020). The number of live and dead cells present after 24 h was monitored using imaging with the Incucyte Live-Cell Analysis System (Sartorius).

### Stable isotope pulse-chase labeling and lipid extraction

Cells were seeded at an initial density of 200,000-500,000 cells per well in a six-well plate in 2 ml of DMEM medium. After incubating for 24 h, cells were washed three times with 2 ml of PBS and then incubated in the indicated media and drug conditions. For 18:2(n-6) isotope labeling experiments, [U-^13^C]-18:2(n-6) (Cambridge Isotope Laboratories, CLM-6855-PK) was supplemented as bovine serum albumin-fatty acid (BSA-FA) conjugates. Stock solutions of 0.8 mM BSA and 0.8 mM BSA/5 mM [U-^13^C]-18:2(n-6) were prepared as described by the Seahorse Bioscience protocol for the XF Analyzer. Solutions were filter-sterilized before addition to cell culture media. For pulse-chase labeling, cells were first treated with 25 µM BSA-[U-^13^C]-18:2(n-6) in DMEM media for 24 h. Cells were then washed three times with 2 ml of PBS before being incubated in the indicated media and drug conditions for a 24 h chase period without label.

For each condition, a parallel plate of cells was scanned with an Incucyte Live-Cell Analysis System (Sartorius) and analyzed for confluence to normalize extraction buffer volumes based on cell number. An empty well was also extracted for a process control. The extraction buffer consisted of chloroform:methanol (containing 25 mg/L of butylated hydroxytoluene (Millipore Sigma, B1378)):0.88% KCl (w/v) at a final ratio of 8:4:3. The final extraction buffer also contained 0.75 µg/ml of norvaline and 0.7 µg/ml of cis-10-heptadecenoic acid as internal standards. For extraction, the medium was aspirated from cells, and cells were rapidly washed in ice-cold saline three times. The saline was aspirated, and methanol:0.88% KCl (w/v) (4:3 v/v) was added. Cells were scraped on ice, and the extract was transferred to 1.5 ml Eppendorf tubes (Dot Scientific, RN1700-GMT) before adding chloroform (Supelco, 1.02444). The resulting extracts were vortexed for 10 min and centrifuged at maximum speed (17000 x g) for 10 min. Lipids (organic fraction) were transferred to glass vials (Supelco, 29651-U) and dried under nitrogen gas for further analysis.

The 3-phase lipid extraction (3PLE) was based on a published protocol^17^. 400 µl of hexane (Supelco, 1.03701), 400 µl of methyl acetate (Sigma Aldrich, 186325), 300 µl of acetonitrile (Fisher Chemical, A9554), and 400 µl of water (Fisher Chemical, W64) were added to the dried lipid pellet. After vortexing, the extracts were centrifuged at maximum speed (3500 rpm) for 5 min, resulting in the separation of three distinct phases. 300 µl of the top hexane phase was collected as the non-polar lipid phase in a glass vial. Non-polar lipids were extracted a second time with another 300 µl of hexane. The polar lipid fraction was obtained by collecting 300 µl of the middle methyl acetate phase into a glass vial. All final extracts were dried under nitrogen gas for further analysis.

### GC-MS analysis of fatty acids

The fatty acid data in Fig. 1A and Fig. 5C were analyzed as fatty acid methyl esters (FAMEs) by GC-MS, as previously described^11^. For all other analyses, fatty acids were analyzed as pentafluorobenzyl-fatty acid (PFB-FA) derivatives. Fatty acids were saponified from dried lipid pellets by adding 800 µl of 90% methanol/0.3 M KOH, vortexing, and incubating at 80°C for 60 min. Each sample was then neutralized with 80 µl of formic acid (Supelco, FX0440). Fatty acids were extracted twice with 800 µl of hexane and dried under nitrogen gas. To derivatize, fatty acid pellets were incubated with 100 µl of 10% pentafluorobenzyl bromide (Sigma Aldrich, 90257) in acetonitrile and 100 µl of 10% N,N-diisopropylethylamine (Sigma Aldrich, D125806) in acetonitrile at room temperature for 30 min. PFB-FA derivatives were dried under nitrogen gas and resuspended in 50 µl of hexane for GC-MS analysis.

GC-MS was conducted with a TRACE TR-FAME column (ThermoFisher, 260M154P) installed in a Thermo Scientific TRACE 1600 gas chromatograph coupled to a Thermo ISQ 7610 mass spectrometer. Helium was used as the carrier gas at a constant flow of 1.8 ml/min. One microliter of sample was injected at 250°C at a 4:1 split (for total lipid extracts) or splitless mode (for 3PLE extracts). After injection, the GC oven was held at 100°C for 0.5 min, increased to 200°C at 40°C/min, held at 200°C for 1 min, increased to 250°C at 5°C/min, and held at 250°C for 11 min. The MS system operated under negative chemical ionization mode with methane gas at a flow rate of 1.25 ml/min, and the MS transfer line and ion source were held at 255°C and 200°C, respectively. The detector was used in scanning mode with an ion range of 150-500 *m/z*. Total ion counts were determined by integrating appropriate ion fragments for each PFB-FA^35^ using Skyline software^36^. Metabolite data were normalized to the internal standard and background-corrected using a process blank sample. Mass isotopologue distributions were corrected for natural abundance using IsoCorrectoR^37^.

### LC-MS analysis of lipids

All samples, blanks, and QC sample pools were resuspended in 50µL of LC/MS grade acetonitrile (A955, Fisher) and LC/MS grade isopropanol (A461, Fisher) 50:50 (v/v), pulse vortexed, and sonicated for 5 min in a room temperature water bath prior to LC-MS/MS analysis. Samples were analyzed with a Vanquish dual pump liquid chromatography system coupled to an Orbitrap ID-X (Thermo Fisher Scientific) using a H-ESI source in both positive and negative mode. All samples were injected at 2 μL and lipids separating with a reversed-phase chromatography using an Accucore C30 column (2.6 μm, 2.1mm × 150mm, 27826-152130, Thermo) combined with an Accucore guard cartridge (2.6 μm, 2.1 mm × 10 mm, 27826-012105, Thermo). Mobile phase A consisted of 60% LC/MS grade acetonitrile (A955, Fisher). Mobile phase B consisted of 90% LC/MS grade isopropanol (A461, Fisher) and 8% LC/MS grade acetonitrile. Both mobile phases contained 10mM ammonium formate (70221, Sigma) and 0.1% LC/MS grade formic acid (A11710X1-AMP, Fisher). Column temperature was kept at 50 °C, flow rate was held at 0.4 mL/min, and the chromatography gradient was as follows: 0-1 min held at 25% B,1-3 min from 25% B to 40% B, 3-19 min from 40% B to 75% B, 19-20.5 min from 75% B to 90% B, 20.5-28 min from 90% B to 95% B, 28-28.1 min from 95% B to 100% B, and 28.1-30 min held at 100% B. A 30 minute re-equilibration went as follows: 0-10 min held at 100% B and 0.2 mL/min, 10-15 min from 100% B to 50% B and held at 0.2 mL/min, 15-20 min held at 50% B and 0.2 mL/min, 20-25 min from 50% B to 25% B and held at 0.2 mL/min, 25-26 min held at 25% B and ramped from 0.2 mL/min to 0.4 mL/min, and 26-30 min held at 25% B and 0.4 mL/min. Each sample was injected twice, once for ESI-positive and once for ESI-negative mode.

Mass spectrometer parameters were: source voltage 3250V for positive mode and −2800V for negative mode, sheath gas 40, aux gas 10, sweep gas 1, ion transfer tube temperature 300°C, and vaporizer temperature 275°C. Full scan data was collected using the orbitrap with scan range of 200-1700 m/z at a resolution of 240,000 for profiling samples and 500,000 for U^13^C6-Glucose traced samples. Primary fragmentation (MS2) was induced in the orbitrap with assisted HCD collision energies at 15, 30, 45, 75, 110%, CID collision energy was fixed at 35%, and resolution was at 15,000. Secondary fragmentation (MS3) was induced in the ion trap with rapid scan rate and CID collision energy fixed at 35% for 3 scans. Lipid identifications were assigned from ddMS3 files using LipidSearch (v5.0, Thermo), which was then used to generate a transition list for peak picking and integration in Skyline (v23.1). For tracing studies, the transition list from LipidSearch was expanded using an in-house script to include every possible carbon isotopologue.

### Quantitative real-time PCR

Total RNA was isolated using the Quick-RNA MiniPrep Plus (Zymo Research, R1057) according to the manufacturer’s protocol. Reverse transcription was performed using iScript gDNA Clear cDNA Synthesis Kit (Bio-Rad, 1725035). Quantitative real-time PCR was performed using the PowerUp SYBR™ Green Master Mix (ThermoFisher, A25741) and Bio-Rad C1000 Touch Thermal Cycler. AGPS primer: sense, 5’-GGAGGCTGCGGGTTCTCTC-3’, antisense, 5’-CCATTTCATAACTTCTTGCCGCT-3’; ATGL primer: sense, 5’-CAATGTCTGCAGCGGTTTCA-3’, antisense, 5’-CCATCCACGTAGCGCACC-3’; ELOVL2 primer: sense, 5’-CGCTGCGGATCATGGAACAT-3’, antisense, 5’-ACCACCCTCTGACTCGAGAA-3’; ELOVL5 primer: sense, 5’-CATTCTCTTGCCGGGGGATT-3’, antisense, 5’-TGCTCCCTTTTGGACTCACA-3’; FADS1 primer: sense, 5’-GTGGCTAGTGATCGACCGTAA-3’, antisense, 5’-GCCACAAAGGGATCCGTGG-3’; FADS2 primer: sense, 5’-GACCACGGCAAGAACTCAAAG-3’, antisense, 5’-GAGGGTAGGAATCCAGCCATT-3’; FAR1 primer: sense, 5’-TGGTATTCCGGAGTTAATAGACCA-3’, antisense, 5’-CGTCTGAAGGCCTGTTCGAG-3’. Quantification of mRNA expression was calculated by the ΔCT method with *18S* rRNA as the reference gene.

### Immunoblotting

Cells were lysed in radioimmunoprecipitation assay (RIPA) buffer (Thermo Scientific, 89900) supplemented with Halt Protease and Phosphatase Inhibitor Cocktail (ThermoFisher, 78440) for 30 min at 4°C. Cell extracts were pre-cleared by centrifugation at maximum speed for 15 min at 4°C, and protein concentration was measured with the Pierce BCA Protein Assay Kit (Thermo Scientific, 23225). Lysates were resolved on SDS-PAGE and transferred electrophoretically to 0.2 µm nitrocellulose membranes (Bio-Rad, 1620112) at 100 V for 60 min. The blots were blocked in Tris-buffered saline buffer (TBST; 10 mmol/L Tris-HCl, pH 8, 150 mmol/L NaCl, and 0.2% Tween-20) containing 5% (w/v) nonfat dry milk for 30 min, and then incubated with the specific primary antibody diluted in blocking buffer at 4°C overnight. Membranes were washed four times in TBST and incubated with HRP-conjugated secondary antibody for 1 h at room temperature. Membranes were washed three times and developed using SuperSignal West Femto Maximum Sensitivity Substrate (Thermo Scientific, 34096). Antibodies were used as follows: ATGL (Cell Signaling Technology 2138, 1:1000), AGPS (Thermo Fisher 56398, 1:1000), Vinculin (Cell Signaling Technology 13901, clone E1E9V, 1:1000), β-actin (Cell Signaling Technology 3700, 1:1000), anti-mouse IgG HRP-linked secondary antibody (Cell Signaling Technology 7076, 1:5000), and anti-rabbit IgG HRP-linked secondary antibody (Cell Signaling Technology 7074, 1:5000).

### Generation of *ATGL*, *FADS2*, and *AGPS* knockout cells by CRISPR-Cas9

sgRNA sequences were designed using CRISPick^38,39^ and cloned into pSpCas9(BB)-2A-GFP (PX458) (Addgene, #48138). A549 cells were transfected with these vectors using Lipofectamine 2000 transfection reagent (Invitrogen, 11668027) according to the manufacturer’s protocol. After 48 hours, GFP-positive cells were sorted into single cells with a BD FACSymphony S6 cell sorter and grown up as single-cell clones. *ATGL* and *AGPS* knockouts were confirmed by immunoblotting, and *FADS2* knockouts were confirmed by next-generation amplicon sequencing (Genewiz, Amplicon-EZ) and analysis using CRISPResso2^40^. sgRNA sequences: *ATGL* exon 5, CGAGAGTGACATCTGTCCGC; *ATGL* exon 3, GATAGCCATGAGCATGCCAG; *FADS2* exon 1, AAGGGAGGGAACCAGGGCGA; *AGPS* exon 1, CAAGGCGGTAGCCATGGCGG; *AGPS* exon 4, ACAGAAGGAGGTGTATCACT.

### Statistics and reproducibility

Sample sizes, reproducibility, and statistical tests used for each figure are denoted in the figure legends. All graphs were generated using GraphPad Prism 9.

## Data availability

All data generated and analyzed during this study are included in this published article. Correspondence and requests for materials should be addressed to Evan C. Lien.

## SUPPLEMENTARY FIGURE LEGENDS

**Figure S1.**
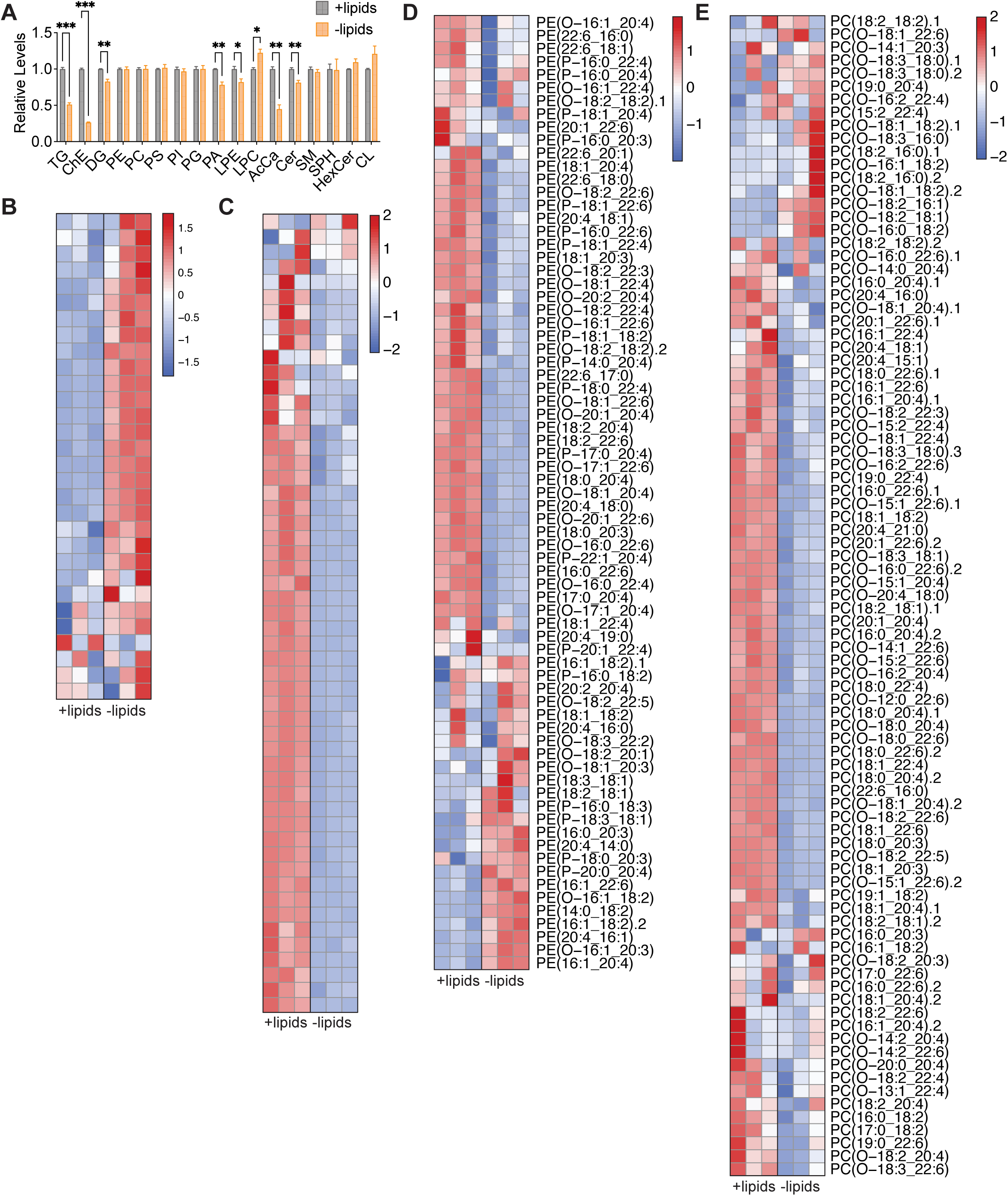
Lipid limitation decreases levels of storage lipids and increases levels of PUFA-containing ether PEs and PCs. HeLa cells were cultured in lipid-depleted versus lipid-replete media for 24 h prior to extracting lipids for analysis by LC-MS. **A,** Relative levels of complex lipid classes. **B,** Heat map showing relative levels of phospholipids containing two MUFA side chains. **C,** Heat map showing relative levels of TGs containing at least one PUFA. **D,** Heat map showing relative levels of PEs containing at least one PUFA. **E,** Heat map showing relative levels of PCs containing at least one PUFA. TG, triglyceride; ChE, cholesterol ester; DG, diglyceride; PE, phosphatidylethanolamine; PC, phosphatidylcholine; PS, phosphatidylserine; PI; phosphatidylinositol; PG, phosphatidylglycerol; LPE, lysophosphatidylethanolamine; LPC, lysophosphatidylcholine; Cer, ceramide; SM, sphingomyelin; SPH, sphingosine; HexCer, hexosylceramide; CL, cardiolipin. Data are presented as mean ± s.e.m; *n* = 3 biologically independent replicates. Comparisons were made using a two-tailed Student’s t-test (**A**). \**P<0.05*, \*\**P<0.01*, \*\*\**P<0.001*.

**Figure S2.**
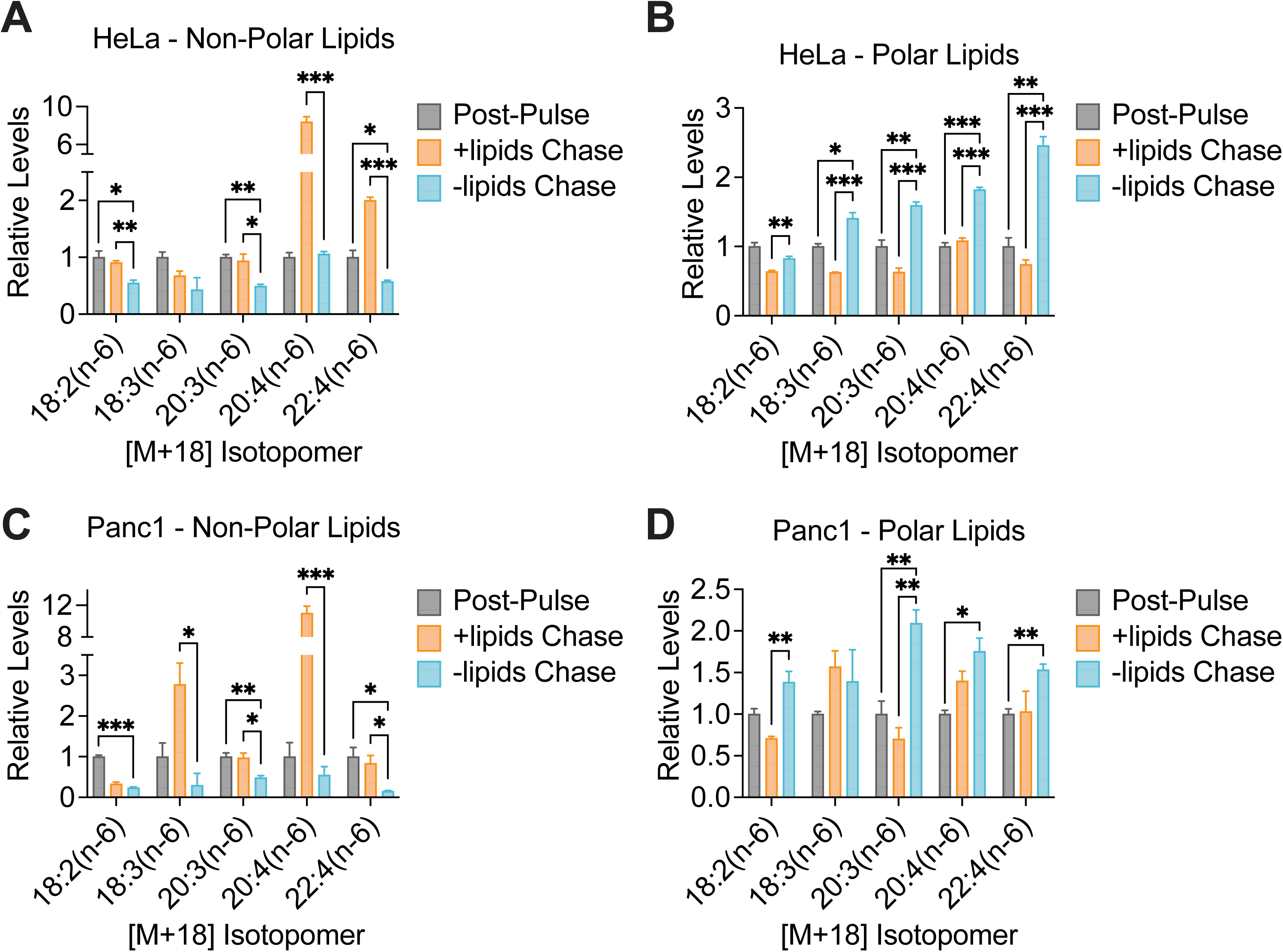
Lipid limitation induces the trafficking of PUFAs from triglycerides into phospholipids. Cells were pulse labeled with [U-^13^C]-18:2(n-6), followed by a chase period prior to lipid extraction and analysis, as shown in Fig. 3A. **A, B,** Relative levels of [M+18] labeled PUFAs in the non-polar (**A**) or polar (**B**) lipid fractions from HeLa cells post-pulse and after the chase period into lipid-depleted versus lipid-replete media. **C, D,** Relative levels of [M+18] labeled PUFAs in the non-polar (**A**) or polar (**B**) lipid fractions from Panc1 cells post-pulse and after the chase period into lipid-depleted versus lipid-replete media. Data are presented as mean ± s.e.m; *n* = 3 biologically independent replicates. Comparisons were made using a two-tailed Student’s t-test (**A-D**). \**P<0.05*, \*\**P<0.01*, \*\*\**P<0.001*.

**Figure S3.**
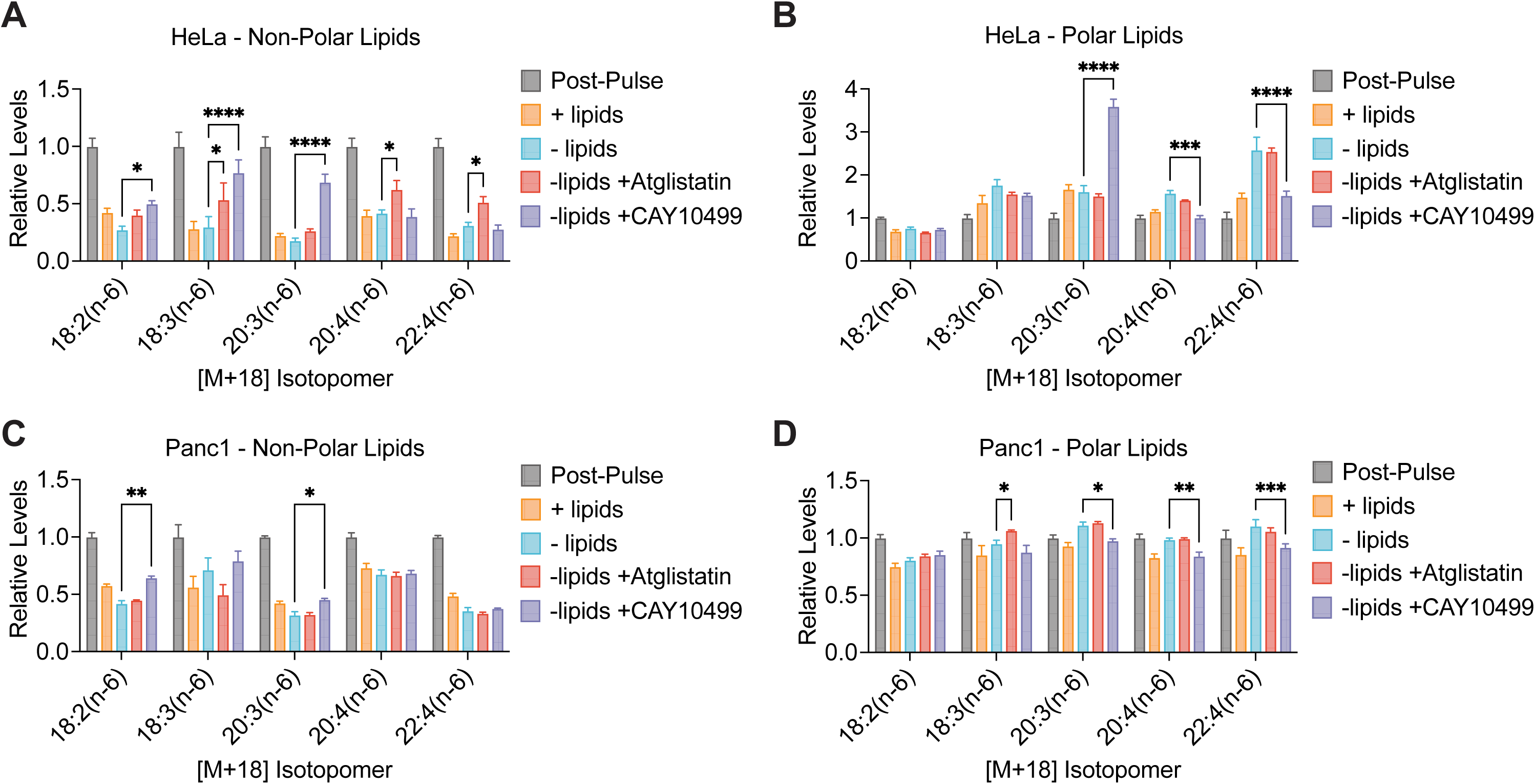
Inhibiting triglyceride lipolysis prevents the accumulation of PUFAs in phospholipids in lipid-starved cancer cells. Cells were pulse labeled with [U-^13^C]-18:2(n-6), followed by a chase period prior to lipid extraction and analysis, as shown in Fig. 3A. **A, B,** Relative levels of [M+18] labeled PUFAs in the non-polar (**A**) or polar (**B**) lipid fractions from HeLa cells post-pulse and after the chase period into lipid-replete media or lipid-depleted media, with or without 10 µM Atglistatin or 5 µM CAY10499. **C, D,** Relative levels of [M+18] labeled PUFAs in the non-polar (**C**) or polar (**D**) lipid fractions from Panc1 cells post-pulse and after the chase period into lipid-replete media or lipid-depleted media, with or without 10 µM Atglistatin or 5 µM CAY10499. Data are presented as mean ± s.e.m; *n* = 3 biologically independent replicates. Comparisons were made using two-way ANOVA (**A-D**). \**P<0.05*, \*\**P<0.01*, \*\*\**P<0.001*.

**Figure S4.**
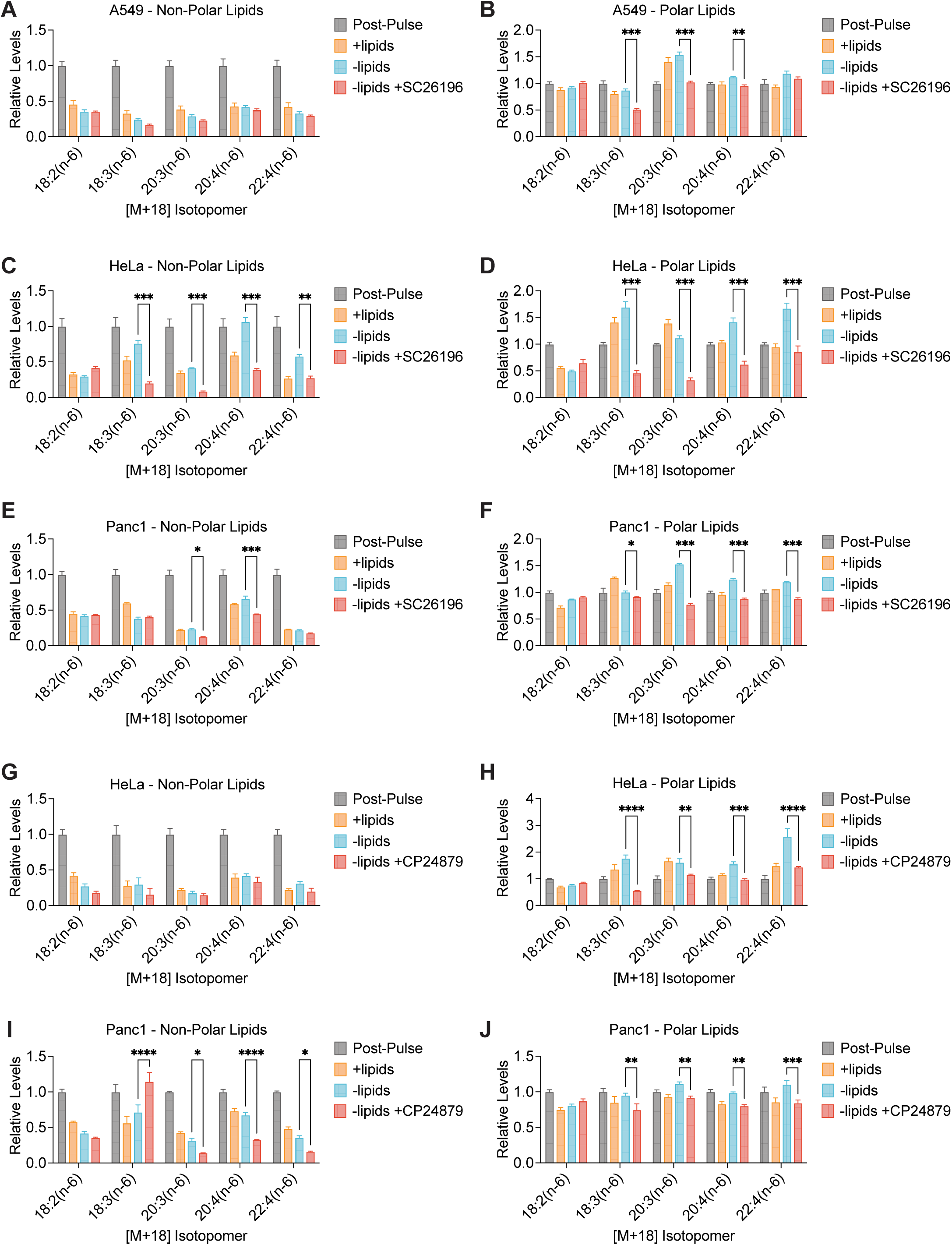
Inhibiting PUFA metabolism prevents the accumulation of PUFAs in phospholipids in lipid-starved cancer cells. Cells were pulse labeled with [U-^13^C]-18:2(n-6), followed by a chase period prior to lipid extraction and analysis, as shown in Fig. 3A. **A-F,** Relative levels of [M+18] labeled PUFAs in the non-polar (**A, C, E**) or polar (**B, D, F**) lipid fractions from A549 (**A, B**), HeLa (**C, D**), and Panc1 (**I, J**) cells post-pulse and after the chase period into lipid-replete media or lipid-depleted media, with or without 5 µM SC26196. **G-J,** Relative levels of [M+18] labeled PUFAs in the non-polar (**G, I**) or polar (**H, J**) lipid fractions from HeLa (**G, H**) and Panc1 (**I, J**) cells post-pulse and after the chase period into lipid-replete media or lipid-depleted media, with or without 10 µM CP24879. Data are presented as mean ± s.e.m; *n* = 3 biologically independent replicates. Comparisons were made using two-way ANOVA (**A-J**). \**P<0.05*, \*\**P<0.01*, \*\*\**P<0.001*.

**Figure S5.**
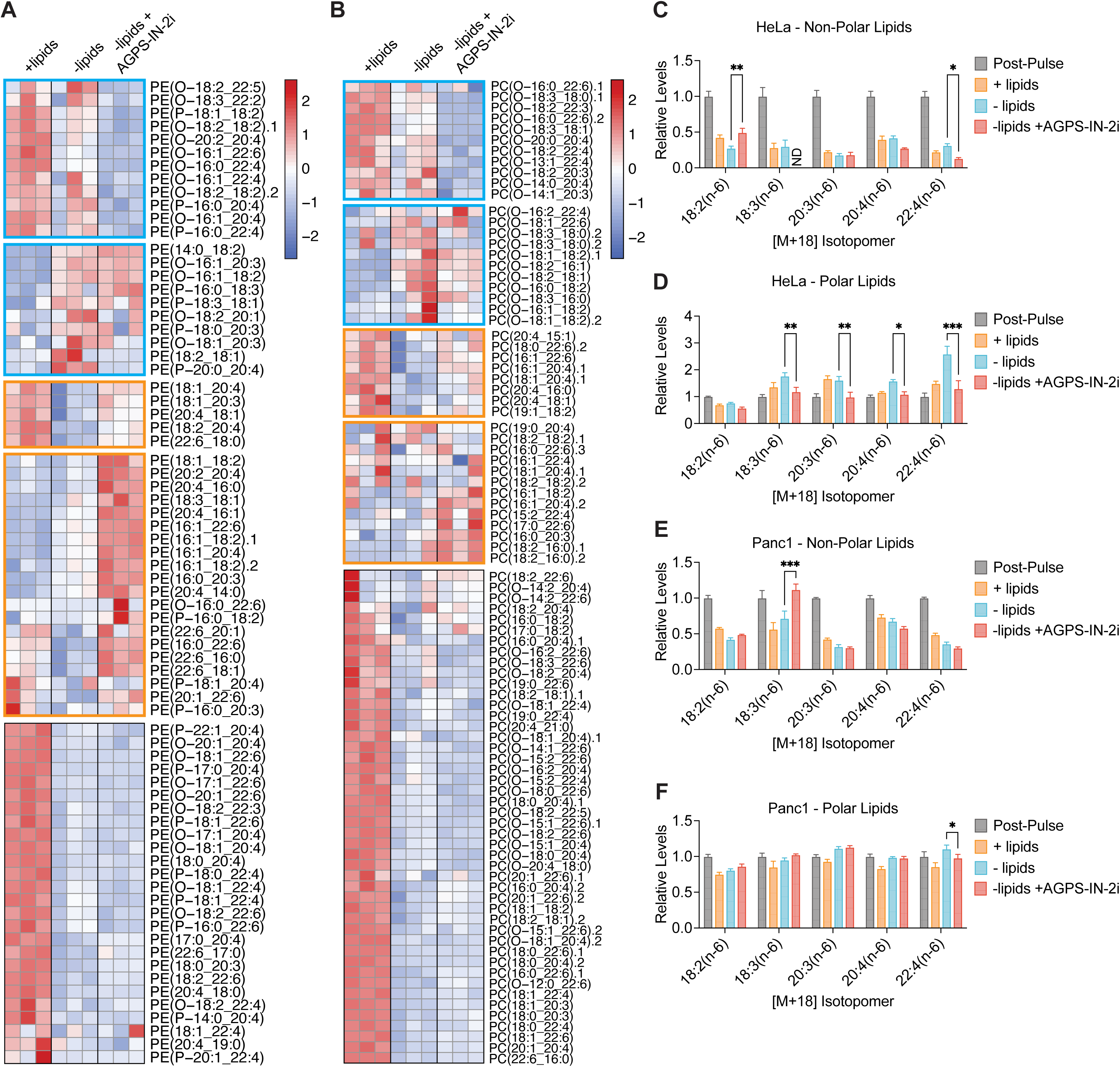
Inhibiting ether lipid synthesis prevents the accumulation of PUFAs in ether phospholipids in lipid-starved cancer cells. **A, B,** Heat map showing relative levels of Pes (**A**) or PCs (**B**) containing at least one PUFA in HeLa cells cultured in lipid-replete or lipid-depleted media, with or without 150 µM AGPS-IN-2i, for 24 h. Blue boxes highlight ether phospholipid species that decrease upon AGPS-IN-2i treatment. Orange boxes highlight diacyl phospholipid species that increase upon AGPS-IN-2i treatment. **C, D,** Cells were pulse labeled with [U-^13^C]-18:2(n-6), followed by a chase period prior to lipid extraction and analysis, as shown in Fig. 3A. Relative levels of [M+18] labeled PUFAs in the non-polar (**C**) or polar (**D**) lipid fractions from HeLa cells post-pulse and after the chase period into lipid-replete media or lipid-depleted media, with or without 150 µM AGPS-IN-2i. **E, F,** Cells were pulse labeled with [U-^13^C]-18:2(n-6), followed by a chase period prior to lipid extraction and analysis, as shown in Fig. 3A. Relative levels of [M+18] labeled PUFAs in the non-polar (**E**) or polar (**F**) lipid fractions from Panc1 cells post-pulse and after the chase period into lipid-replete media or lipid-depleted media, with or without 150 µM AGPS-IN-2i. Data are presented as mean ± s.e.m; *n* = 3 biologically independent replicates. Comparisons were made using two-way ANOVA (**C-F**). \**P<0.05*, \*\**P<0.01*, \*\*\**P<0.001*.

